# Hyperactive Ras disrupts cell size control and a key step in cell cycle entry in budding yeast

**DOI:** 10.1101/2023.03.06.531344

**Authors:** Jerry T. DeWitt, Jennifer C. Chinwuba, Douglas R. Kellogg

## Abstract

Severe defects in cell size are a nearly universal feature of cancer cells. However, the underlying causes are unknown. A previous study suggested that a hyperactive mutant of yeast Ras (*ras2^G19V^*) that is analogous to the human Ras oncogene causes cell size defects, which could provide clues to how oncogenes influence cell size. However, the mechanisms by which *ras2^G19V^* influences cell size are unknown. Here, we found that *ras2^G19V^* inhibits a critical step in cell cycle entry, in which an early G1 phase cyclin induces transcription of late G1 phase cyclins. Thus, *ras2^G19V^*drives overexpression of the early G1 phase cyclin Cln3, yet Cln3 fails to induce normal transcription of late G1 phase cyclins, leading to delayed cell cycle entry and increased cell size. *ras2^G19V^* influences transcription of late G1 cyclins via a poorly understood step in which Cln3 inactivates the Whi5 transcriptional repressor. Previous studies found that Ras relays signals via protein kinase A (PKA) in yeast; however, *ras2^G19V^*appears to influence G1 phase cyclin expression via novel PKA-independent signaling mechanisms. Together, the data define new mechanisms by which hyperactive Ras influences cell cycle entry and cell size in yeast. Expression of G1 phase cyclins is also strongly influenced by mammalian Ras via mechanisms that remain unclear. Therefore, further analysis of PKA-independent Ras signaling in yeast could lead to discovery of conserved mechanisms by which Ras family members control expression of G1 phase cyclins.

## Introduction

Cells within the human body range in size over several orders of magnitude. However, cells of a particular type maintain a constant average size. Thus, cell growth must be tightly controlled to ensure that cells attain and maintain an appropriate cell size (Liu et al., 2022; Jorgensen and Tyers, 2004; Turner et al., 2012). At the simplest level, the size and shape of a cell must be the outcome of conserved mechanisms that determine the extent, location, and timing of cell growth. In dividing cells, maintenance of a specific cell size is ensured by mechanisms that link cell cycle progression to cell growth. These mechanisms are modulated by nutrients such that the threshold amount of growth required for cell cycle progression is reduced in poor nutrients, which leads to a reduction in cell size (Kellogg and Levin, 2022). The mechanisms that control cell growth and size remain poorly understood.

Previous studies in budding yeast suggested that a Ras signaling network is required for normal control of cell size. Yeast Ras is encoded by a pair of redundant paralogs referred to as *RAS1* and *RAS2*. Cells lacking either paralog are viable but loss of both is lethal (Toda et al., 1985). The functions of yeast Ras are best understood in the context of a signaling network that is activated by glucose. High glucose activates Ras1/2, which then activate adenylate cyclase to produce cAMP. The cAMP binds Bcy1, an inhibitory subunit for the yeast homolog of cAMP-dependent protein kinase (PKA) (Matsumoto et al., 1983; Toda et al., 1987). Binding of cAMP to Bcy1 causes it to dissociate from PKA, leading to release of active PKA that initiates a signaling network with pervasive effects on control of cell growth and metabolism (Robinson et al., 1987; Zaman et al., 2009). A hyperactive allele of RAS2 (*ras2^G19V^*) that is analogous to oncogenic alleles of mammalian Ras was found to cause an increase in cell size in diploid cells that also contain a mutant allele of *CDC25*, the budding yeast Ras-GEF (Broek et al., 1987; Baroni et al., 1989); however, the effects of hyperactive *ras2^G19V^*on cell size have not been tested in a wild type background. Deletion of *IRA2*, a budding yeast Ras-GAP that is conserved in mammals, also causes an increase in cell size (Jorgensen et al., 2002). Furthermore, a weakly constitutive allele of PKA in a *bcy1*Δ background causes a failure in nutrient modulation of cell size (Tokiwa et al., 1994). Together, these observations suggest that a Ras-PKA signaling axis influences cell size and plays a role in nutrient modulation of cell size; however, the mechanisms are poorly understood. Furthermore, little is known about yeast Ras signaling beyond the PKA pathway and it remains possible that Ras influences cell size via signaling pathways that are distinct from the canonical Ras-PKA pathway.

Ras proteins are highly conserved and mammalian Ras homologs serve as master regulators of growth, proliferation, metabolism and survival (Cazzanelli et al., 2018; Weiss, 2020). Ras family members are amongst the most frequently mutated oncogenic drivers; it is thought that between 25-30% of all human cancers have oncogenic mutations in one or more Ras genes (Hobbs et al., 2016). A potential role for Ras family members in controlling cell size is intriguing, since severe defects in cell size homeostasis are broadly linked to cancer (Asadullah et al., 2021; Asa, 2019; Hoda et al., 2018; Gothwal et al., 2021; Brimo et al., 2013; Sandlin et al., 2022). For example, most cancers are associated with greater heterogeneity of cell size and shape, as well as dramatically altered nuclear-to-cytoplasmic volume ratios. Defects in cell size and shape have long been used by pathologists to diagnose cancer, and increased heterogeneity of cell size is associated with poor prognosis (Asadullah et al., 2021). The size defects of cancer cells must be caused, either directly or indirectly, by oncogenic signals. However, the mechanisms by which oncogenic signals influence cell growth and size are largely unknown, and it is unclear whether the size defects of cancer cells are a direct consequence of primary oncogenic signals or a secondary consequence of mutations that accumulate during evolution of cancer cells. The fact that diverse cancers show common defects in cell size suggests the possibility that diverse oncogenic signals converge on common conserved pathways for size control.

Since it is unknown how oncogenic signals influence cell size in human cells, or how hyperactive Ras influences cell size in yeast, it is possible that oncogenic Ras influences cell size via mechanisms that are conserved from yeast to humans. Thus, analysis of how hyperactive Ras influences cell size in yeast could provide new clues to the functions of oncogenic Ras in human cells. Here, we carried out new analyses of the effects of constitutively active Ras in budding yeast, utilizing modern methods that allow rapid inducible expression of *ras2^G19V^*at endogenous levels in otherwise wild type cells, which allowed us to discern the immediate consequences of *ras2^G19V^*activity during the cell cycle. We found that hyperactive Ras strongly influences expression of G1 phase cyclins, potentially via signaling mechanisms that are independent of the canonical Ras-PKA signaling axis. Ras also influences expression of G1 phase cyclins in mammalian cells, potentially via signaling mechanisms that are distinct from the canonical MAP kinase pathway that is downstream of Ras (Muise-Helmericks et al., 1998; Pruitt et al., 2000; Pruitt and Der, 2001; Coleman et al., 2004; Kerkhoff and Rapp, 1998). Together, these observations raise the possibility that Ras controls G1 phase cyclin expression via novel unexplored pathways that may be conserved from yeast to humans.

## Results

### Hyperactive Ras increases cell size in budding yeast

Previous experiments examined the effects of hyperactive Ras expressed from the endogenous promoter on cell size in a mutant background (Baroni et al., 1989). We therefore started by analyzing the effects of hyperactive Ras on cell size in an otherwise wild type background. A hyperactive version of *RAS2* can be generated by mutating glycine 19 to a valine (*ras2^G19V^*), which is analogous to oncogenic versions of mammalian Ras. We generated a strain that expresses *ras2^G19V^* from the endogenous promoter and measured cell size with a Coulter Channelyzer. The *ras2^G19V^* allele caused an increase in cell size (**Figure 1A**). The effects of *ras2^G19V^* were slightly stronger in a *ras1Δ* background (**Figure 1–figure supplement 1A**)

**Figure 1.**
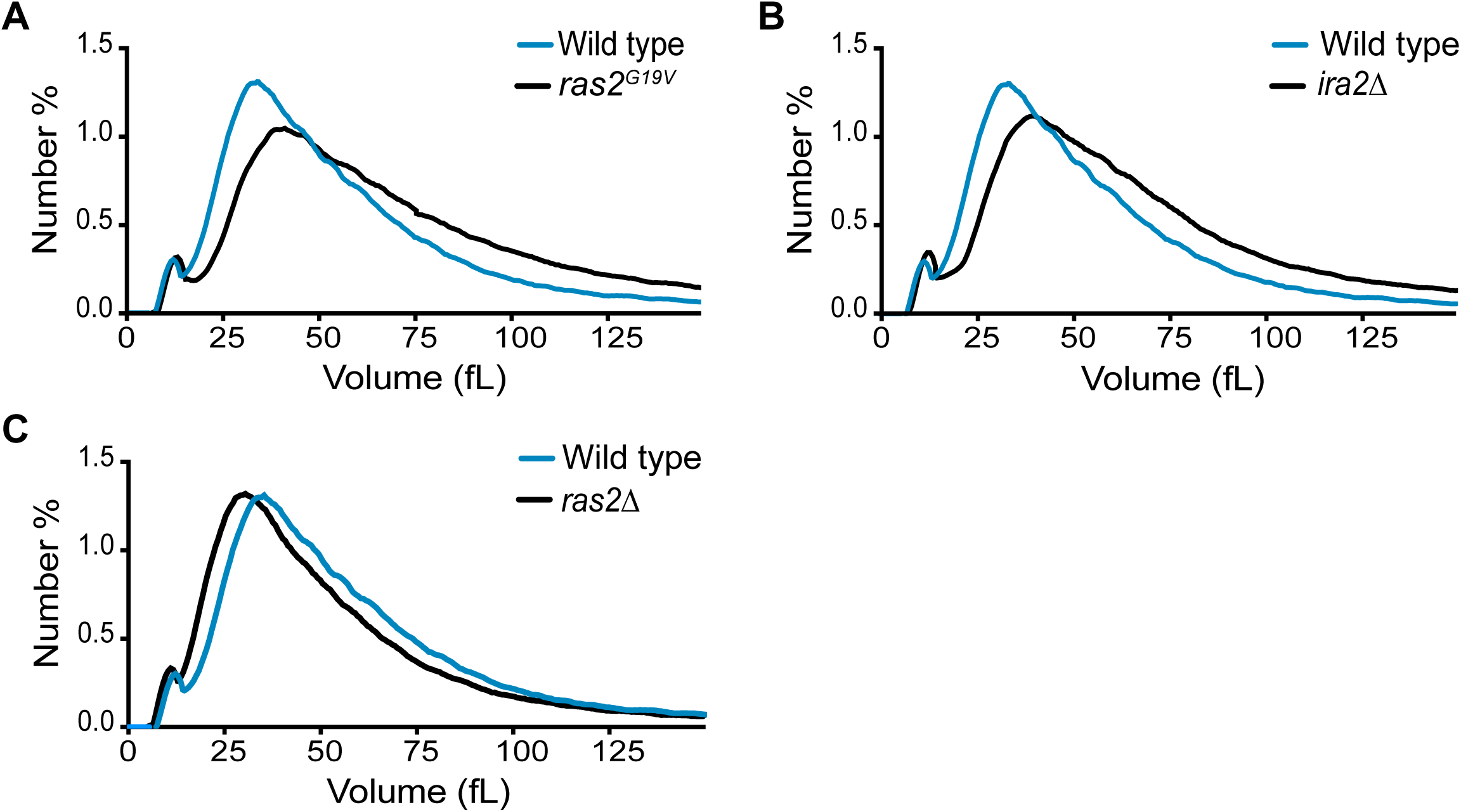
Hyperactive Ras increases cell size. (A) Wild type and *ras2^G19V^* yeast cells were grown to log phase in YPD and cell size was measured using a Coulter counter. (B) Wild type and *ira2Δ* budding yeast cells were grown to log phase in YPD and cell size was measured using a Coulter counter. (C) Wild type and *ras2*Δ budding yeast cells were grown to log phase in YPD and cell size was measured using a Coulter counter.

As an independent means of generating hyperactive Ras, we deleted one of the GAPs that contribute to inactivation of Ras. GAPs for Ras are encoded by two partially redundant paralogs referred to as *IRA1* and *IRA2*. Deletion of *IRA2* causes cells to proliferate more slowly, whereas deletion of *IRA1* does not have an obvious phenotype. Deletion of both genes is lethal. We found that *ira2Δ* caused an increase in cell size, which provided further evidence that hyperactive Ras influences cell size (**Figure 1B**). A previous genome-wide search for gene deletions that influence cell size also found that *ira2Δ* causes increased cell size (Jorgensen et al., 2002). We next tested the effects of loss of function of Ras1 and Ras2. While *ras1Δ* did not affect cell size, *ras2Δ* caused a modest decrease (**Figure 1C and Figure 1–figure supplement 1B**). Together, these results establish that Ras activity influences cell size.

### An inducible and titratable system for expression of *ras2^G19V^* in yeast

A previous study found that *ras2^G19V^*causes decreased viability (Toda et al., 1985), and we found that *ras2^G19V^* cells show a large reduction in the rate of proliferation and rapidly accumulate suppressor mutations. The effects of hyperactive Ras on cell size could therefore be an indirect consequence of long-term adaptation to constitutive Ras activity. To circumvent this problem, we utilized a previously developed estradiol-inducible promoter to achieve tight temporal and titratable control of *RAS* gene expression in a nutrient independent manner (Ottoz et al., 2014). In brief, cells were engineered to express a fusion protein that includes the bacterial LexA DNA-binding domain, the human estrogen receptor (ER), and a transcriptional activation domain (AD). In addition, a promoter containing two LexA binding sites was integrated in front of the wild type endogenous *RAS2* coding sequence or in front of *ras2^G19V^*. In this context, addition of ß-estradiol drives transcription of *RAS2* or *ras2^G19V^* **(Figure 2A)**.

**Figure 2.**
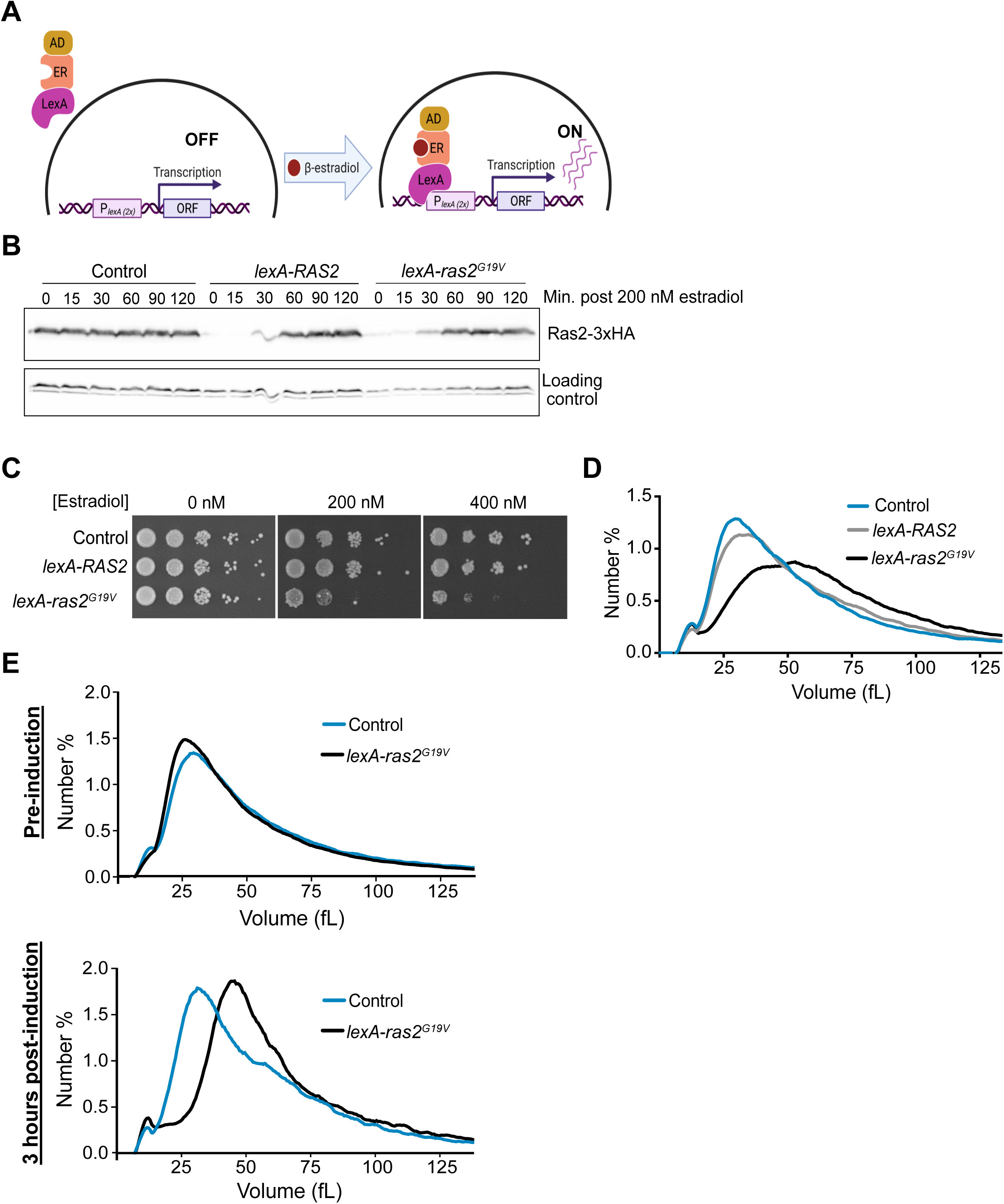
An inducible and titratable expression system for expression of *ras2^G19V^* in yeast. (A) A diagram of the LexA-ER-AD system for estradiol-dependent gene expression. (B) Estradiol was added to 200 nM and time points were collected at the indicated intervals to measure timing for peak levels of Ras2 or ras2^G19V^ protein expression relative to endogenous levels. (C) Serial dilutions of the indicated strains were spotted onto YPD medium containing the indicated concentration of estradiol. (D) Cells were grown overnight to log phase in YPD + 100 nM estradiol and cell size was measured using a Coulter counter. (E) Cells were grown overnight to log phase in YPD and expression of *ras2^G19V^* was induced with 100 nM estradiol for 3 hours. Cell size was measured using a Coulter counter.

To determine the time required to reach peak protein expression, we created strains that express *RAS2-3xHA* and *ras2^G19V^-3xHA* from the *lexA* promoter (*lexA-RAS2-3xHA* and *lexA-ras2^G19V^-3xHA)*. Peak protein expression was reached within 60-90 minutes after addition of estradiol (**Figure 2B**). Moreover, we found that 200 nM estradiol induced expression of *lexA-RAS2-3xHA* and *lexA-ras2^G19V^-3xHA* at protein levels similar to *RAS2-3xHA* expressed from the endogenous promoter.

For analysis of *ras2^G19V^*phenotypes, we utilized untagged versions of *RAS2* (*lexA-RAS2* and *lexA-ras2^G19V^*). We first tested the effects of inducing expression of *ras2^G19V^* on cell proliferation. Serial dilutions of cells containing *lexA-ras2^G19V^* and control cells were plated on media containing increasing concentrations of estradiol. Expression of *ras2^G19V^*at endogenous levels (100-200 nM estradiol) caused a large reduction in the rate of proliferation (**Figure 2C**). Expression at higher levels of *ras2^G19V^*with 400 nM estradiol was nearly lethal.

We next tested whether expression of *ras2^G19V^* from the *lexA* promoter causes an increase in cell size. Prolonged expression of *ras2^G19V^* for 12 hours caused a large increase in cell size (**Figure 2D**). The increase in cell size was detectable within 3 hours of inducing expression, which suggests that it is a rapid and direct consequence of ras2^G19V^ protein expression (**Figure 2E**).

### Expression of *ras2^G19V^* influences cell cycle progression and cell size

We next set out to learn more about how *ras2^G19V^* influences cell size. To do this, we first analyzed how *ras2^G19V^* influences the relationship between cell growth and cell cycle progression. Previous studies analyzed the effects of hyperactive Ras on the cell cycle indirectly by manipulating cAMP levels to mimic Ras-dependent signals that control PKA, or by deleting the BCY1 gene, which leads to a high level of constitutive PKA signaling (Matsumoto et al., 1983; Tokiwa et al., 1994; Toda et al., 1987; Baroni et al., 1994; Hall, 1998; Mizunuma et al., 2013). Together, these studies suggested a link between cAMP-dependent signaling and regulation of G1 cyclin expression; however, in some cases the studies reached conflicting conclusions. For example, two studies found that cAMP-dependent signaling stimulates production of *CLN3* mRNA (Mizunuma et al., 2013; Tokiwa et al., 1994) whereas other studies observed no effect (Hall, 1998; Hubler et al., 1993). Similarly, several studies found that cAMP-dependent signaling increases transcription of *CLN1* and *CLN2* (Hall, 1998; Hubler et al., 1993) whereas two other studies reported a decrease in transcription (Tokiwa et al., 1994; Baroni et al., 1994). Differences in results could be due to technical differences in how the experiments were carried out.

Previous investigations were limited by the tools available at the time and were unable to analyze the immediate effects of Ras-dependent signaling on the cell cycle. Moreover, although production of cAMP is a well-established output of Ras-dependent signaling in budding yeast, it is possible that Ras has targets beyond cAMP production. An additional limitation of previous studies is that they did not analyze how cAMP-dependent signals influence G1 cyclin protein expression during the cell cycle.

Here, we directly examined how expression of *lexA-ras2^G19V^* influences cell cycle progression, cell growth, and cyclin expression in synchronized cells. Cells were arrested in G1 phase with mating pheromone and expression of *RAS2* or *ras2^G19V^*from the *lexA* promoter was induced prior to release from the arrest. Wild type cells that express *RAS2* from the endogenous promoter were included as a control. Since bud emergence marks cell cycle entry, we first analyzed the percentage of cells undergoing bud emergence as a function of time. Expression of *lexA-ras2^G19V^* caused a prolonged delay in bud emergence (**Figure 3A**). We also analyzed cell size as a function of time with a Coulter Channelyzer (**Figure 3B**) and plotted bud emergence as a function of cell size (**Figure 3C**). *ras2^G19V^* caused cells to undergo bud emergence at a larger cell size. These data are consistent with a previous report that increased levels of cAMP inhibit cell cycle entry and cause increased cell size at the G1/S transition (Tokiwa et al., 1994).

**Figure 3.**
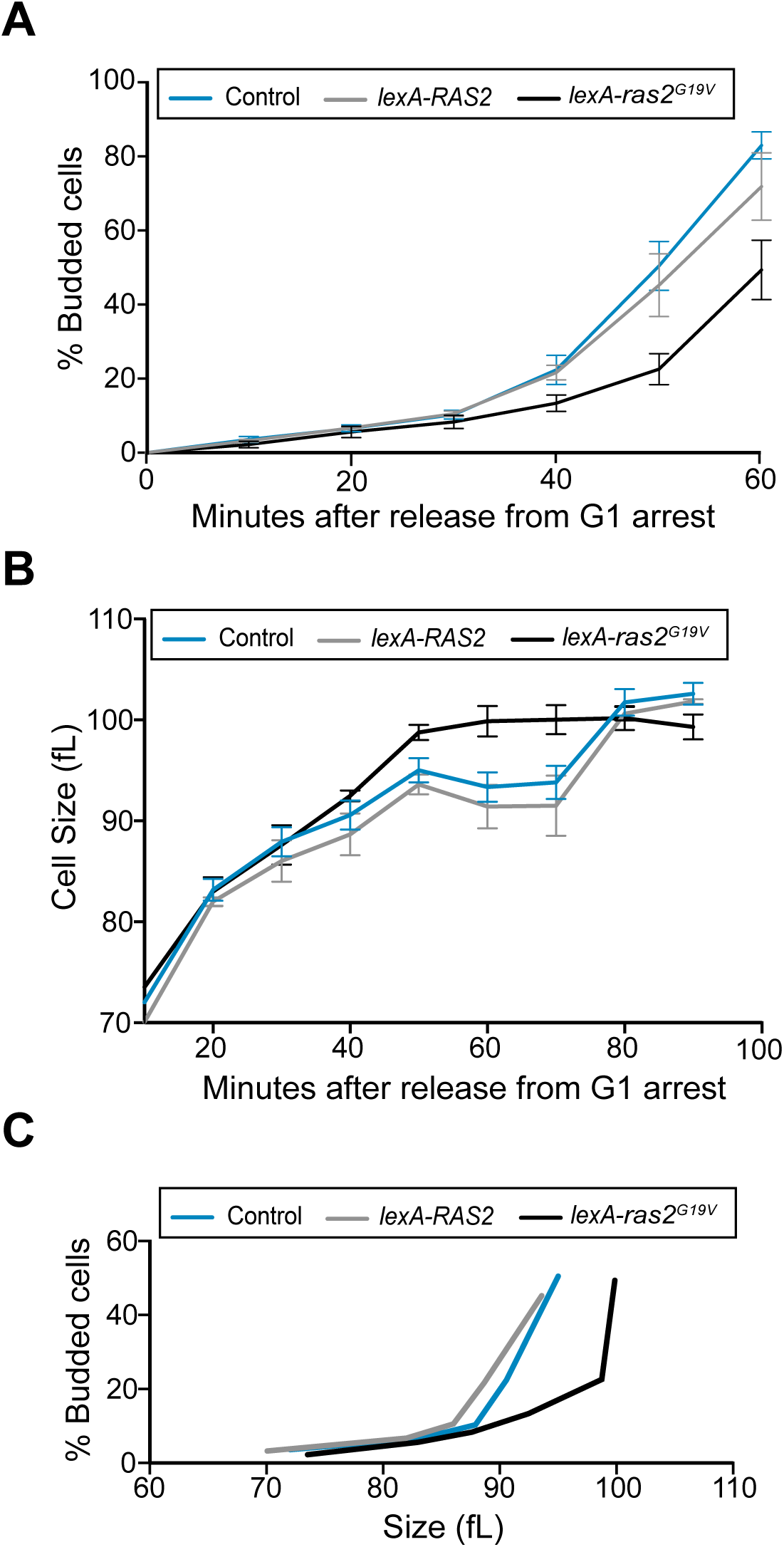
Expression of *ras2^G19V^* influences cell size and delays cell cycle entry. (A-C) Cells were grown to log-phase overnight in YPD and were then arrested in G1 phase with alpha factor for 3 hours at 30°C. Cells were treated with 200 nM estradiol beginning 1.5 hours before release from the arrest. (A) The percentage of budded cells as a function of time. (B) Median cell size was measured at 10 min intervals using a Coulter Counter and plotted as a function of time. (C) The percentage of budded cells as a function of cell size. Error bars represent SEM of three biological replicates.

### Expression of *ras2^G19V^* blocks a key step in cell cycle entry

We next analyzed the effects of *ras2^G19V^*on expression of G1 phase cyclins. In budding yeast, a cyclin called *CLN3* is expressed in early G1 phase and accumulates gradually (Zapata et al., 2014; Lucena et al., 2018; Landry et al., 2012). Accumulation of Cln3 protein is correlated with growth and appears to be dependent upon cell growth, which suggests that it could be a readout of the extent of growth (Zapata et al., 2014; Lucena et al., 2018; Sommer et al., 2021). Cln3 eventually triggers transcription of a redundant pair of late G1 cyclin paralogs called *CLN1* and *CLN2* (Tyers et al., 1993; Dirick et al., 1995; Stuart and Wittenberg, 1995). Expression of late G1 cyclins is the key molecular event that drives cell cycle entry. To assay accumulation of both early and late G1 cyclins, we used a strain that contains Cln3-6xHA and Cln2-13xMyc. Wild type, *lexA-RAS2* and *lexA-ras2^G19V^* cells were released from a G1 arrest and the behavior of Cln3-6xHA and Cln2-13xMyc were assayed by western blot during the cell cycle. *ras2^G19V^*caused a nearly 3-fold increase in the expression of Cln3-6xHA protein in G1 phase, as well as delayed and decreased expression of Cln2-13xMyc (**Figure 4 and Figure 4–figure supplement 1A**). Expression of *lexA-ras2^G19V^* also caused a decrease in Cln2-13Myc levels in asynchronously growing cells (**Figure 4–figure supplement 1B**).

**Figure 4.**
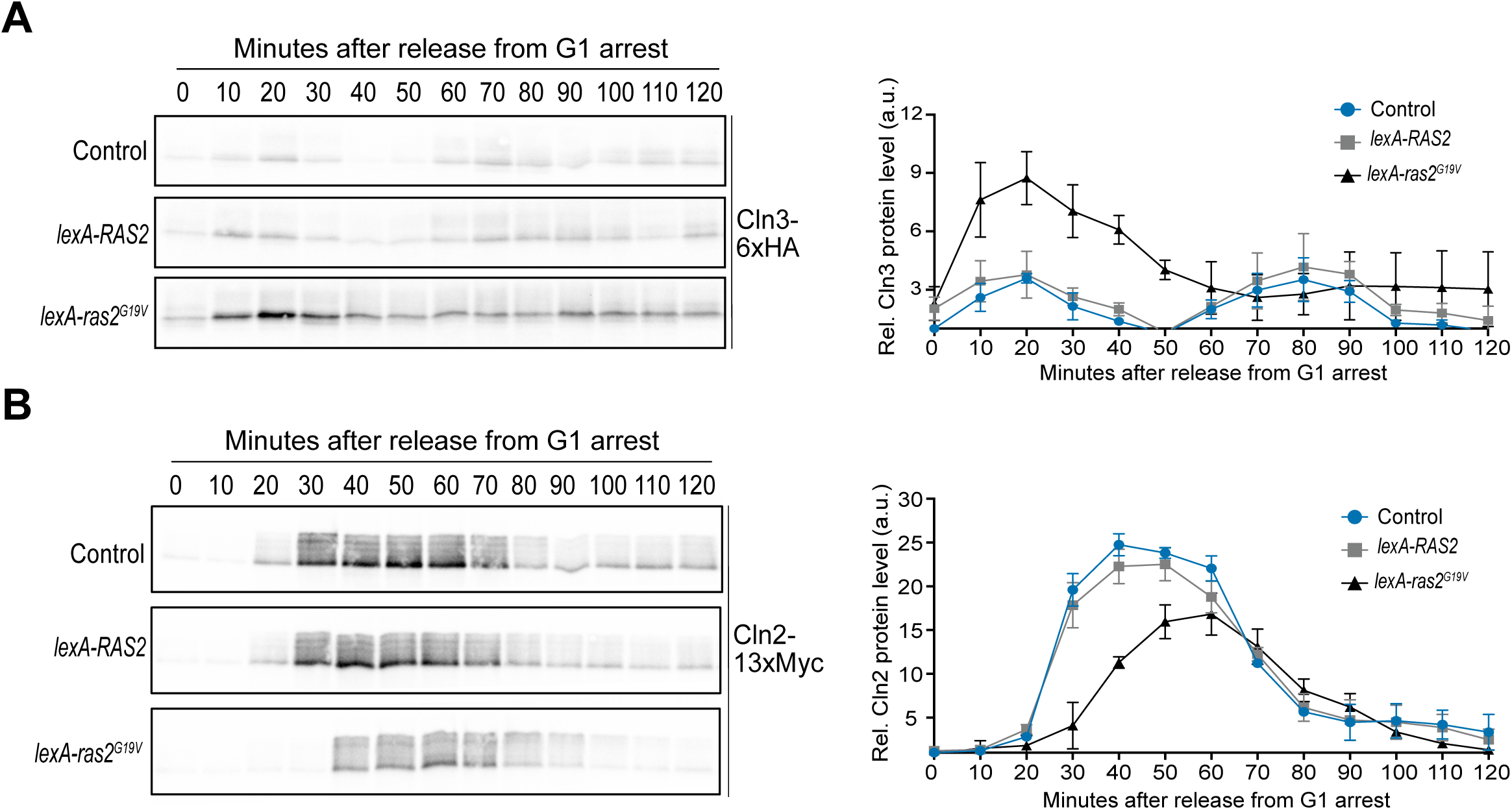
*ras2^G19V^* influences expression of G1 phase cyclins. (A-B) Cells were grown to log-phase overnight in YPD and were then arrested in G1 phase with alpha factor for 3 hours at 30°C. Expression of *RAS2* or *ras2^G19V^*was induced with 200 nM estradiol beginning 1.5 hours before release from the arrest. (A) The levels of Cln3-6xHA protein were analyzed by western blot. (B) The levels of Cln2-13xMyc protein were analyzed by western blot. The representative western blots in (A) and (B) are from the same experiment and protein abundance was quantified relative to levels in wildtype cells at time point 0 after first normalizing to a loading control, see Figure 4–figure supplement 1. Error bars represent SEM of three biological replicates. Loading controls are shown in Figure 4 –figure supplement 1. For both plots, protein abundance was plotted as a ratio over the signal in wild type control cells at t=0 after first normalizing to the loading control.

Previous studies suggested that increased expression of *CLN3* drives premature cell cycle entry (Cross, 1988; Nash et al., 1988; Nasmyth and Dirick, 1991). Thus, the finding that *ras2^G19V^*strongly promotes expression of Cln3, yet inhibits expression of Cln2, was unexpected.

We next used northern blotting to test whether *ras2^G19V^* influences transcription of *CLN2*. *ras2^G19V^* caused a delay in *CLN2* transcription and a reduction of *CLN2* mRNA levels (**Figure 5A**). To test whether *ras2^G19V^* also influences expression of *CLN2* via post-transcriptional mechanisms, we replaced the endogenous *CLN2* promoter with the heterologous *MET25* promoter. In this context, *ras2^G19V^*no longer delayed Cln2 protein expression and caused a small decrease in protein levels, although it was unclear whether the decrease was significant (**Figure 5B**). This observation suggests that *ras2^G19V^*influences *CLN2* expression largely via transcriptional mechanisms.

**Figure 5.**
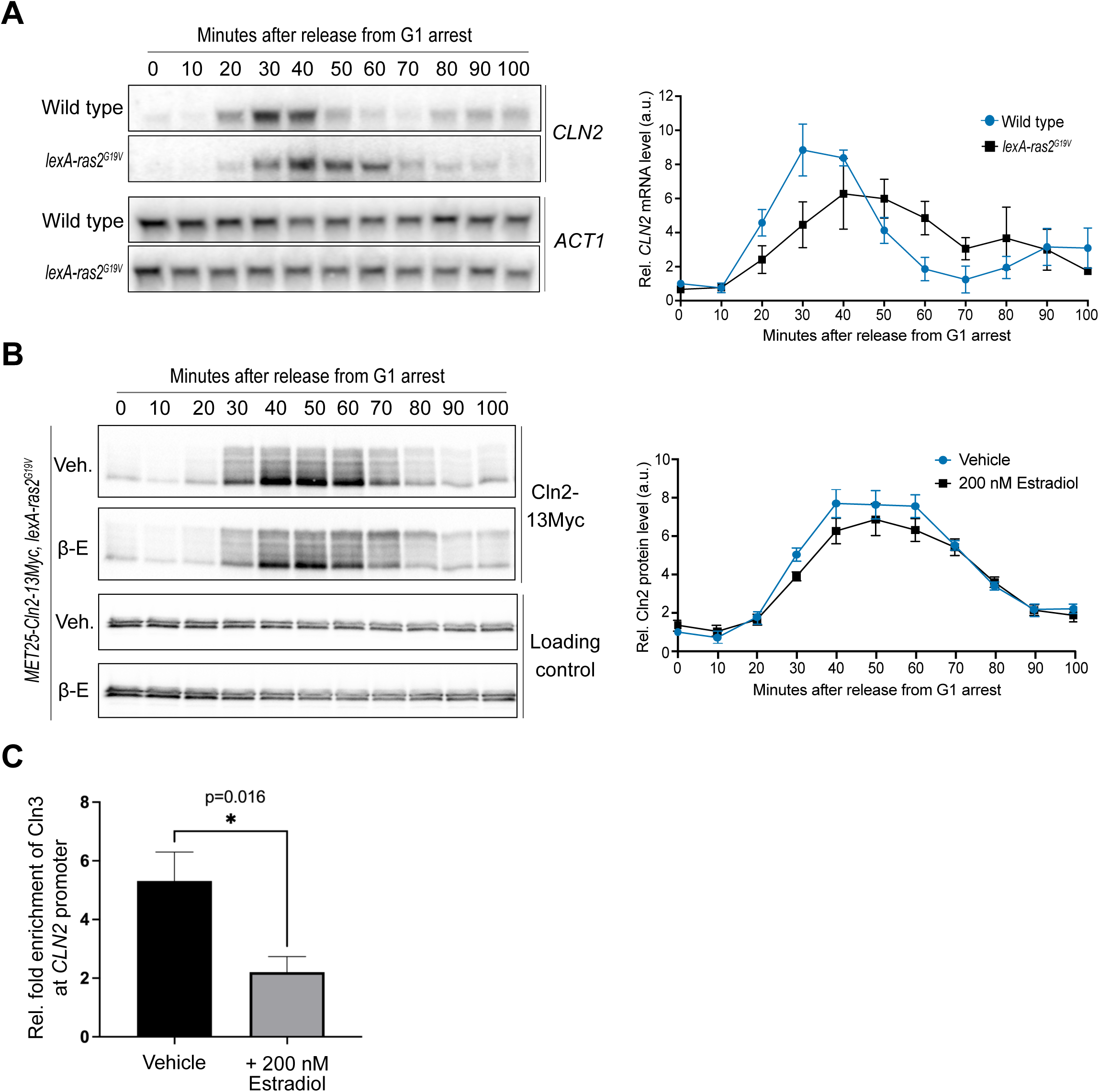
Expression of *ras2^G19V^* modulates CLN2 at the level of transcription. (A-B) Cells were grown to log-phase overnight in YPD and were then arrested in G1 phase with alpha factor for 3 hours at 30°C. Expression of *RAS2* or *ras2^G19V^* was induced with 200 nM estradiol beginning 1.5 hours before release from the arrest. (A) Levels of *CLN2* mRNA were analyzed by northern blot. Levels of the *ACT1* mRNA were analyzed as a loading control. After normalizing to the loading control, the *CLN2* mRNA signal was quantified and plotted as a ratio over the signal at t=0 in the wild control cells. Error bars represent SEM of three biological replicates. (B) Levels of Cln2-13xMyc protein were analyzed by western blot. Protein abundance was plotted as a ratio over the signal in wildtype cells at t=0 after first normalizing to the loading control. Error bars represent SEM of three biological replicates. An anti-Nap1 antibody was used for a loading control. (C) For ChIP experiments, cells were grown to log-phase overnight in YPD and expression of *RAS2* or *ras2^G19V^* was induced with 200 nM estradiol for 3 hours at 30°C. Cln3-6xHA was immunoprecipitated and qPCR was conducted to determine relative fold enrichment of Cln3-6xHA at the *CLN2* promoter. *p = 0.016 by student’s t test.

Induction of late G1 cyclin transcription by Cln3 is thought to be the critical molecular step that initiates cell cycle entry. It is also thought to be the step where cell growth influences cell cycle entry. Thus, the discovery that expression of *ras2^G19V^*promotes high level expression of Cln3, yet fails to induce normal expression of Cln2, suggests that *ras2^G19V^*inhibits a key step in cell cycle entry.

### *ras2^G19V^* causes decreased recruitment of Cln3 to the *CLN2* promoter

Previous work has shown that Cln3 is recruited to promoters controlled by SBF, including the *CLN2* promoter, where it has been proposed to activate Cdk1 to directly phosphorylate RNA polymerase (Kõivomägi et al., 2021; Wang et al., 2009). To test whether *ras2^G19V^* disrupts recruitment of Cln3 to the *CLN2* promoter, we used ChIP to analyze recruitment of Cln3 to *CLN2* promoters in *lexA-ras2^G19V^* cells. We found that *ras2^G19V^* caused a 3-fold reduction in the amount of Cln3 that is recruited to the *CLN2* promoter (**Figure 5C**). The mechanisms by which Cln3 is recruited to the *CLN2* promoter are unknown so we are unable to define the molecular defect that obstructs Cln3 recruitment.

### PKA influences cell size and expression of G1 phase cyclins

The fact that Ras activates PKA suggests that the effects of *ras2^G19V^* on cell size could be mediated by PKA. Furthermore, it has been proposed that PKA can influence expression of G1 phase cyclins via inhibition of Whi3, an RNA binding protein that is thought to bind and inhibit the expression of dozens of mRNAs, including those for *CLN2* and *CLN3* (Wang et al., 2004; Mizunuma et al., 2013; Cai and Futcher, 2013). *WHI3* was originally discovered in a screen for loss of function mutants that cause reduced cell size (Nash et al., 2001). Although one study found no effect of *whi3Δ* on Cln3 protein expression (Garí et al., 2001), a more recent study found that *whi3Δ* causes an increase in the abundance and translation efficiency of the *CLN3* mRNA, which could account for the reduced cell size of *whi3Δ* cells (Cai and Futcher, 2013; Wang et al., 2004). Mutation of a PKA consensus site within Whi3 causes reduced binding of Whi3 to *CLN3* mRNA; however, it remains unknown whether this site is phosphorylated by PKA in vivo (Mizunuma et al., 2013). Together, these observations suggest a model in which *ras2^G19V^*influences G1 phase cyclin levels and cell size via PKA-dependent inhibition of Whi3.

To begin to test this model, we first analyzed the effects of modulating PKA activity on cell size and cell cycle progression. Prior work tested the effects of increased PKA activity by deleting the *BCY1* gene, which encodes the inhibitory subunit for PKA (Matsumoto et al., 1983; Toda et al., 1985, 1987; Guerra et al., 2022). Loss of *BCY1* leads to increased cell size (Baroni et al., 1994; Tokiwa et al., 1994). However, we found that *bcy1Δ* is lethal in the strain background used here (W303). We therefore used an analog-sensitive version of PKA to analyze the effects of decreased PKA activity. PKA in budding yeast is encoded by three redundant genes referred to as *TPK1*, *TPK2* and *TPK3*. A previous study generated cells in which all three *TPK* genes carry mutations that make them sensitive to the adenine analog inhibitor 1NM-PP1 (*pka-as*) (Zaman et al., 2009). If the effects of *ras2^G19V^* are due to hyperactive PKA, then reduced activity of PKA would be expected to cause effects that are opposite to the effects of hyperactive Ras alone.

We found that *pka-as* cells showed substantially reduced cell size in response to a non-lethal dose of inhibitor (50 nM) (**Figure 6A**). Inhibition of PKA had little effect on peak levels of Cln3, although it did prolong the interval of expression of Cln3 in G1 phase **(Figure 6B)**. Expression of Cln2 protein was reduced and delayed by inhibition of PKA, similar to the effects of *ras2^G19V^*.

**Figure 6.**
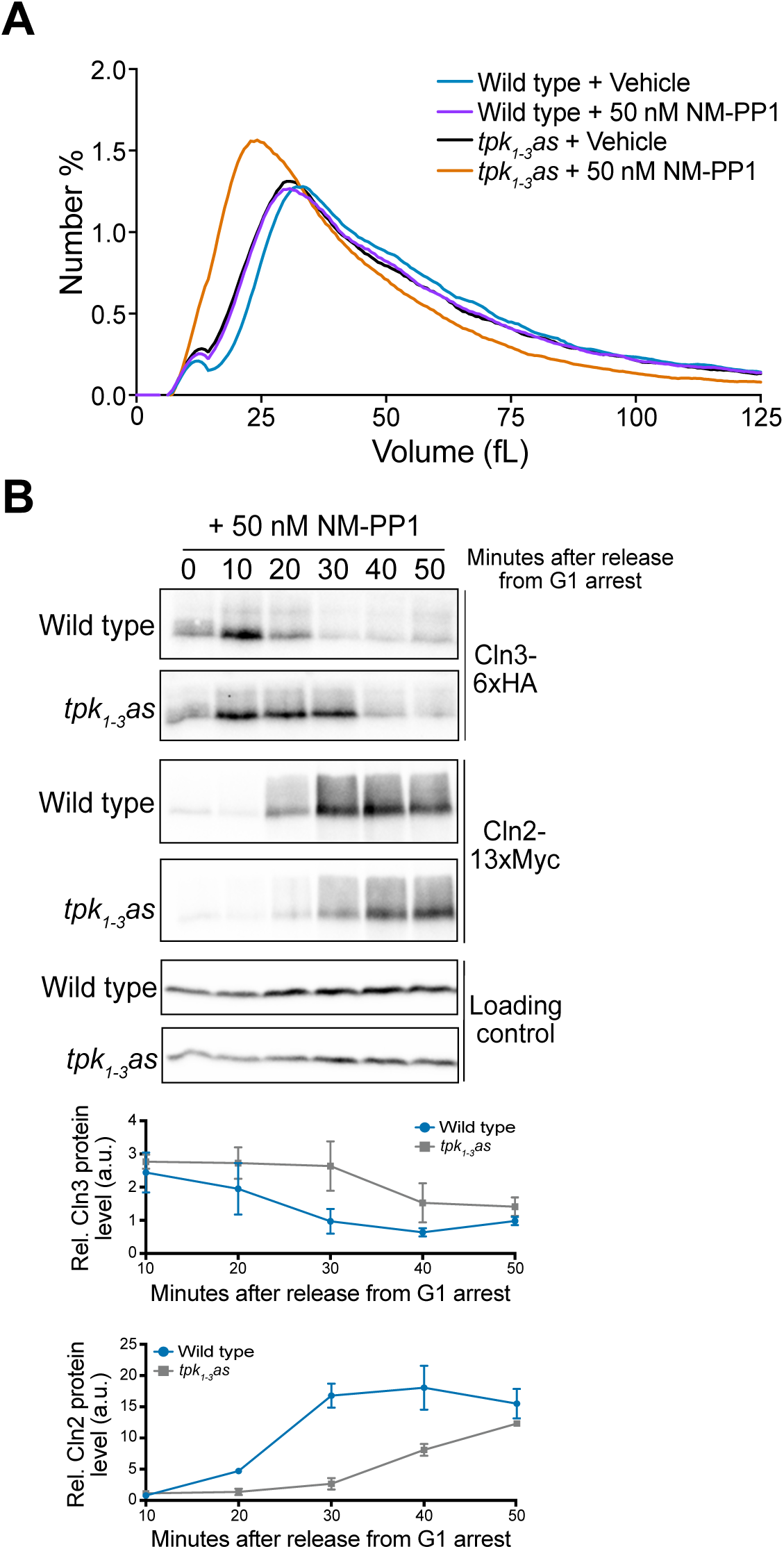
PKA activity influences cell size and expression of G1 phase cyclins. (A) Cells were grown overnight to log phase in YPD + vehicle or 50 nM 1NM-PP1 and cell size was measured using a Coulter Counter. (B) Cells were grown to log-phase overnight in YPD and were then arrested in G1 phase with alpha factor for 3 hours at 30°C. Cells were released from the arrest into YPD containing 100 nM 1NM-PP1. The levels of Cln3-6xHA and Cln2-13xMyc protein were analyzed by western blot. Protein abundance was plotted as a ratio over the signal in wildtype cells at t=0 after first normalizing to the loading control. Error bars represent SEM of three biological replicates. Anti-Nap1 was used for a loading control.

The observed effects of inhibiting PKA are difficult to reconcile with simple models for *ras2^G19V^*functions based on previous studies. The discovery that *ras2^G19V^*and inhibition of PKA have opposite effects on cell size would appear to be consistent with a model in which *ras2^G19V^*drives an increase in cell size via hyperactivation of PKA. However, the discovery that *ras2^G19V^* and inhibition of PKA have similar effects on Cln2 protein levels and that inhibition of PKA has little effect on peak levels of Cln3 suggests that the effects caused by hyperactive Ras signaling cannot be explained solely by changes in the activity of PKA. Rather, the data suggest that hyperactive Ras signaling could be influencing Cln2 levels via PKA-independent mechanisms.

### The effects of *ras2^G19V^* are not due solely to inhibition of Whi3

To investigate the relationship between *ras2^G19V^* signaling and Whi3, we compared cell size, the timing of bud emergence, and expression of Cln3 and Cln2 protein in wildtype, *whi3Δ*, *lexA-ras2^G19V^*, and *lexA-ras2^G19V^ whi3Δ* cells. If *ras2^G19V^* influences expression of Cln3 or Cln2 via PKA-dependent inhibition of Whi3, one would expect *whi3*Δ and *ras2^G19V^* to cause similar effects on expression of Cln3 and Cln2 protein levels.

We found that *whi3*Δ accelerated the timing of bud emergence and caused reduced cell size, as expected (**Figure 7A,B**). Moreover, in contrast to a previous study (Garí et al., 2001) we found that *whi3Δ* caused a substantial increase in Cln3 protein levels (**Figure 7C, Figure supplement 3**). *whi3Δ* also caused a strong increase in the second peak of Cln3 later in the cell cycle (**Figure 7C and Figure 7–figure supplement 1**). Note that alpha factor was added back to the cells to prevent a second cell cycle, so the second peak in Cln3 levels corresponds to the second mitotic peak of Cln3 that has been reported previously (Zapata et al., 2014; Landry et al., 2012). *whi3Δ* also accelerated expression of Cln2 protein, as expected for increased Cln3 protein levels (**Figure 7D and Figure 7–figure supplement 1**).

**Figure 7.**
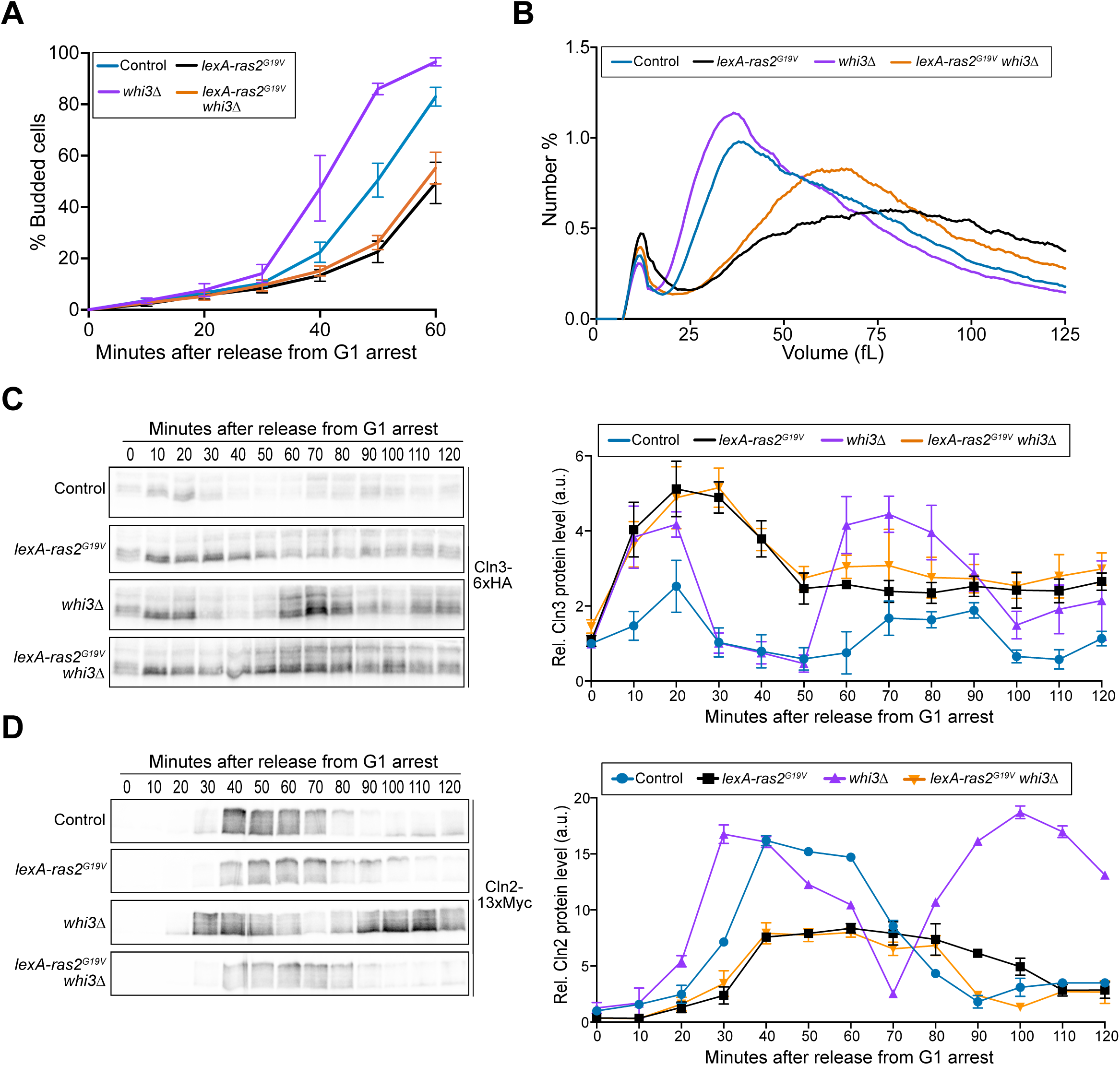
The effects of *ras2^G19V^* are not due solely to inhibition of Whi3. (A-C) Cells were grown to log-phase overnight in YPD and were then arrested in G1 phase with alpha factor for 3 hours at 30°C. Cells were treated with 200 nM estradiol beginning 1.5 hours prior to release. (A) The percentage of budded cells as a function of time. (B) The levels of Cln3-6xHA protein were analyzed by western blot and protein abundance was quantified relative to levels of Cln3-6xHA in wildtype cells at t=0 after first normalizing to a loading control. The loading control is shown in Figure 7-figure supplement 1. Error bars represent SEM of three biological replicates. (C) The levels of Cln2-13xMyc protein were analyzed by western blot and protein abundance was quantified relative to levels of Cln2-13xMyc in wildtype cells at t=0 after first normalizing to a loading control. The loading control is shown in Figure 7-figure supplement 1. Error bars represent SEM of three biological replicates. Error bars represent SEM of three biological replicates. (D) Cells were grown overnight to log phase in YPD + 100 nM estradiol and cell size was measured using a Coulter counter.

The effects of *lexA-ras2^G19V^* expression and *whi3Δ* on Cln3 protein levels were similar, as both caused a substantial increase in Cln3 protein levels. Moreover, the effects of *whi3Δ* and *lexA-ras2^G19V^* on peak Cln3 protein levels were not additive in G1 phase in *lexA-ras2^G19V^ whi3Δ* cells. Thus, it is possible that *ras2^G19V^* drives an increase in Cln3 protein levels via inhibition of Whi3. However, *whi3Δ* and expression of *lexA-ras2^G19V^* caused opposite effects on the timing of bud emergence, cell size, and accumulation of Cln2 protein. Thus, the effects of *ras2^G19V^* cannot be explained solely by inhibition of Whi3.

### The effects of *ras2^G19V^* on late G1 cyclin expression are dependent upon Whi5

Genetic data suggest that Cln3 promotes transcription of *CLN2* at least partly via inhibition of Whi5, which binds and inhibits the SBF transcription factor that promotes *CLN2* transcription (de Bruin et al., 2004; Costanzo et al., 2004). We found that *whi5Δ* largely rescued the delays in Cln2 expression and cell cycle progression caused by *ras2^G19V^***(Figure 8A)**. This observation provides further support for the idea that expression of *lexA-ras2^G19V^*blocks a key step in the mechanisms by which Cln3 initiates transcription of *CLN2*, and it suggests that *ras2^G19V^*may prevent inhibition of Whi5. However, the mechanisms by which Cln3 promotes inhibition of Whi5 are poorly understood (Kõivomägi et al., 2021; Bhaduri et al., 2015).

**Figure 8.**
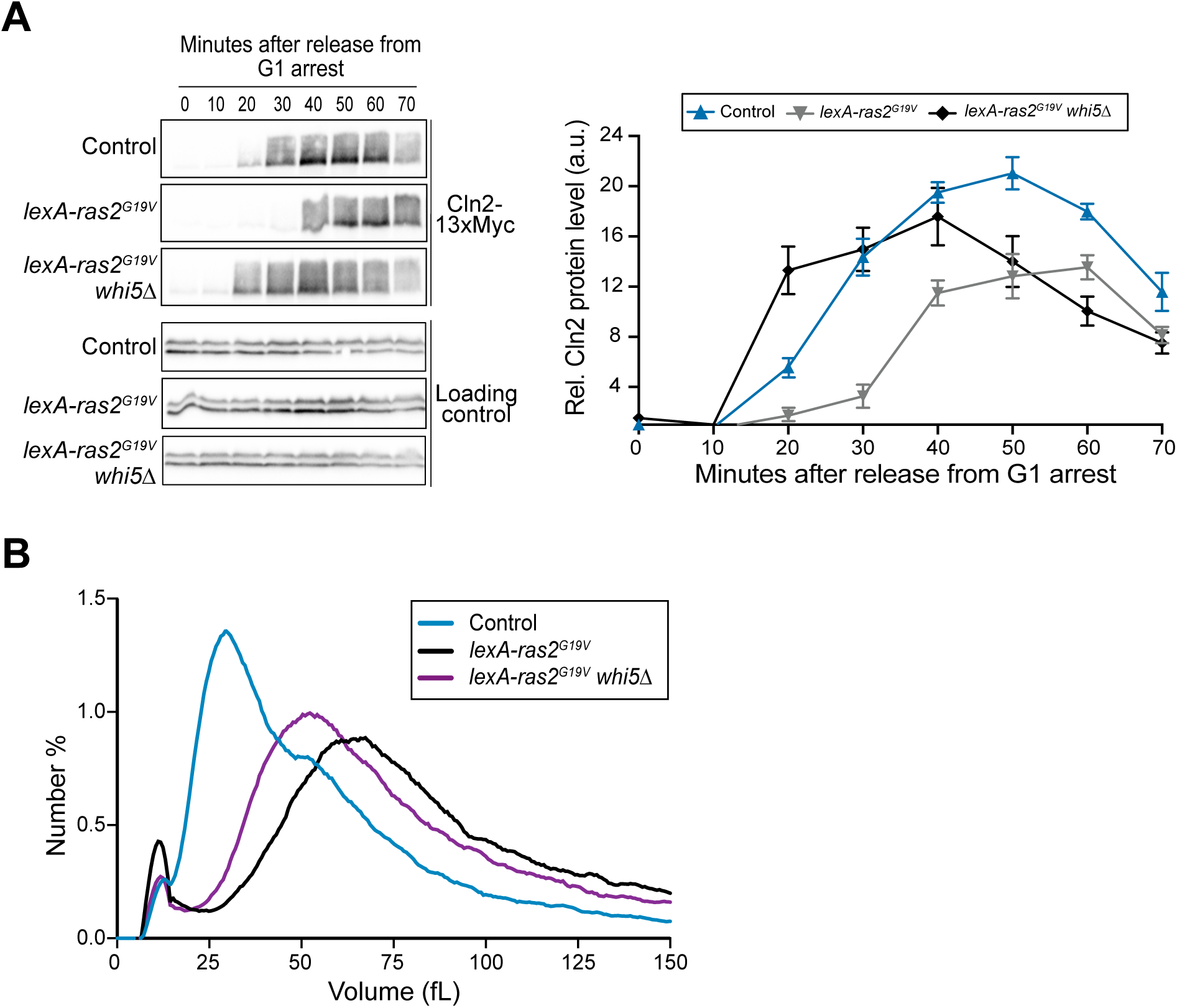
The effects of *ras2^G19V^*on late G1 phase cyclin expression are dependent upon Whi5. (A) Cells were grown to log-phase overnight in YPD and then arrested in G1 phase with alpha factor for 3 hours at 30°C. Cells were treated with 200 nM estradiol beginning 1.5 hours prior to release. The levels of Cln2-13xMyc protein were analyzed by western blot and protein abundance was quantified relative to levels of Cln2-13xMyc in wildtype cells at t=0 after first normalizing to the loading control. Error bars represent SEM of three biological replicates. An anti-Nap1 antibody was used for a loading control. (B) Cells were grown overnight to log phase in YPD + vehicle or 100 nM estradiol and cell size was measured using a Coulter counter.

Deletion of *WHI5* only partially rescued the cell size defects caused by *ras2^G19V^*, which suggests that *ras2^G19V^*influences cell size via mechanisms that operate outside of G1 phase **(Figure 8B)**.

### Cln2 may influence expression of Cln3 via negative feedback

*ras2^G19V^*and inhibition of PKA both caused Cln3 protein to persist for a longer interval in G1 phase (**Figures 4A, 6B, and 7C**). In each context, prolonged expression of Cln3 protein was accompanied by reduced and delayed accumulation of Cln2. A potential explanation for these observations is that Cln2 is required for downregulation of Cln3. This kind of negative feedback regulation has been observed at other stages in the cell cycle. For example, mitotic cyclins repress expression of cyclins that appear earlier in the cell cycle (Amon et al., 1993). To test this idea, we determined whether gain-of or loss-of-function of *CLN1/2* influences expression of Cln3 protein in synchronized cells. We found that over-expression of *CLN2* from the *GAL1* promoter led to substantially lower levels of Cln3 protein during G1 phase, as well as a premature reduction in Cln3 protein levels (**Figure 9A**). Conversely, loss of function of Cln1/2 in *cln1Δ cln2Δ* cells appeared to cause increased Cln3 protein expression during G1 phase (**Figure 9B**). A caveat is that the *cln1Δ cln2Δ* cells did not fully synchronize. Therefore, we also analyzed Cln3 levels in unsynchronized cells, which again appeared to show that loss of CLN1/2 causes an increase in Cln3 protein levels (**Figure 9C**).

**Figure 9.**
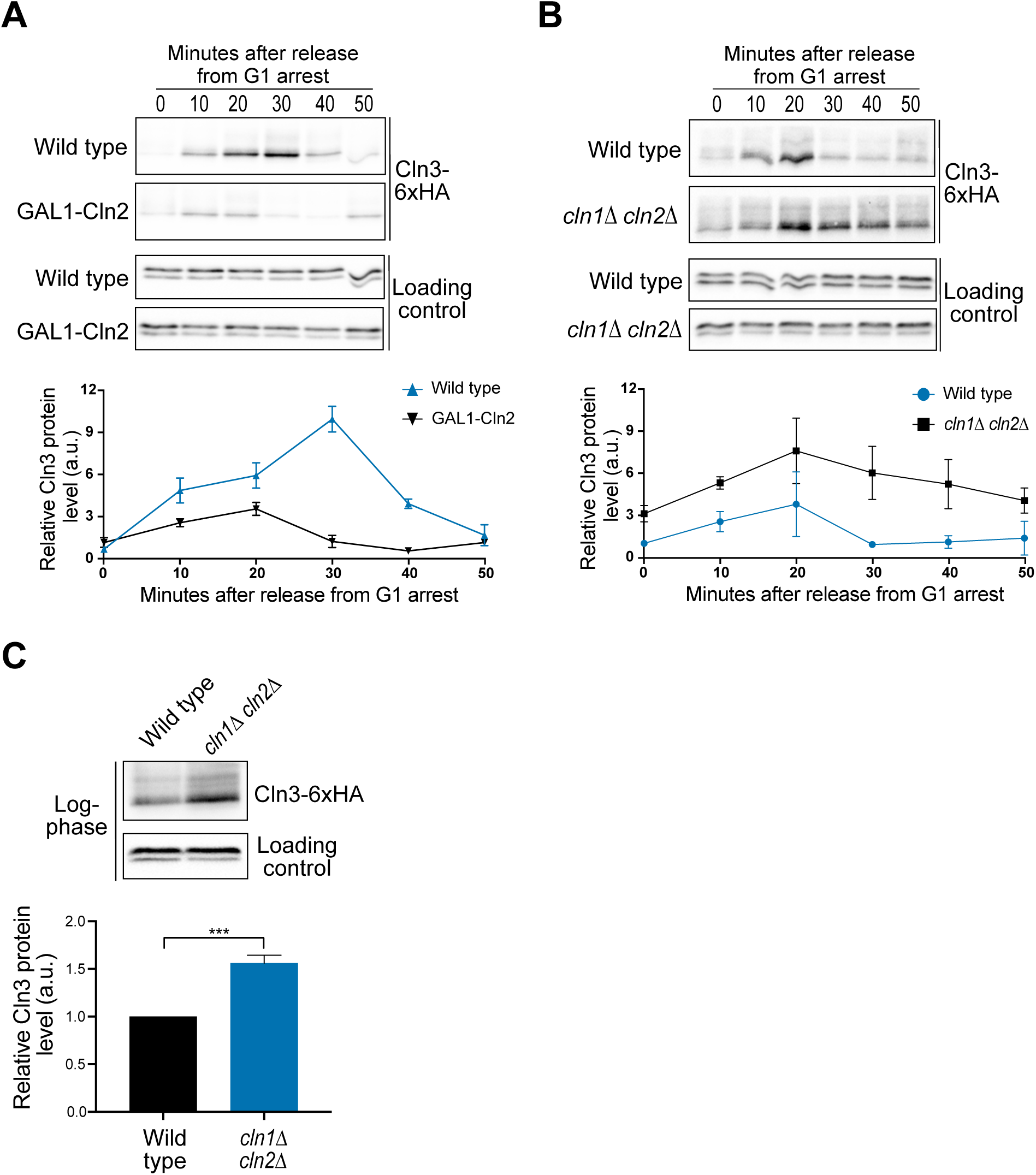
Cln2 imposes negative feedback upon Cln3. (A) Cells were grown to log-phase overnight in YPG/E and were then arrested in G1 phase with alpha factor for 3 hours at 30°C. Expression of GAL1-CLN2 was induced 40 minutes before release by addition of 2% galactose. Cells were washed and released into YEP containing 2% galactose. The levels of Cln3-6xHA protein were analyzed by western blot and protein abundance was quantified relative to Cln3-6xHA levels in wildtype cells at t=0 after first normalizing to the loading control. (B) Cells were grown to log-phase overnight in YPD and were then arrested in G1 phase with alpha factor for 3 hours at 30°C. The levels of Cln3-6xHA protein were analyzed by western blot and protein abundance was quantified relative to Cln3-6xHA levels in wildtype cells at t=0 after first normalizing to the loading control. (C) Cells were grown to log-phase overnight in YPD. Levels of Cln3-6xHA protein were analyzed by western blot and protein abundance was quantified relative to Cln3-6xHA levels in wildtype. (A-C) Error bars represent SEM of three biological replicates. *** p < 0.001 by student’s t test.

## Discussion

### Hyperactive Ras influences cell size and disrupts a critical step in cell cycle entry

Pioneering work carried out over 20 years ago suggested that Ras influences cell size in yeast. However, technical limitations made it difficult to determine whether size defects caused by Ras mutants were an immediate and direct consequence of aberrant Ras activity. Here, we used modern methods to express Ras from an inducible promoter at endogenous levels, which establishes a powerful new system in which to analyze the effects of hyperactive Ras. This allowed us to clearly establish that cell size defects are an immediate and direct consequence of *ras2^G19V^*activity.

Several previous studies suggested that hyperactive Ras influences cell cycle progression and expression of G1 phase cyclins, which could help explain the size defects of *ras2^G19V^* cells. However, these studies examined the effects of hyperactive Ras indirectly by manipulating PKA signaling, and in some cases obtained conflicting results (Matsumoto et al., 1983; Toda et al., 1985, 1987; Tokiwa et al., 1994; Baroni et al., 1994; Hall, 1998; Mizunuma et al., 2013). Here, we directly tested the immediate effects of *ras2^G19V^* expressed from an inducible promoter, which showed that *ras2^G19V^* causes a prolonged delay before bud emergence. Growth continues during the delay so that *ras2^G19V^*cells initiate bud emergence at a substantially larger size than control cells.

To better understand the cause of the G1 phase delay, we assayed expression of early and late G1 phase cyclins and found that *ras2^G19V^* causes a 3-fold increase in Cln3 protein levels and also prolongs Cln3 expression in G1 phase. Increased Cln3 expression would be expected to accelerate and increase expression of Cln2. However, we found that *ras2^G19V^* causes delayed and decreased expression of *CLN2* mRNA and protein. The effects of *ras2^G19V^* on Cln2 protein expression appeared to occur primarily at the level of transcription, since replacing the normal promoter of *CLN2* with a heterologous promoter eliminated much of the effect of *ras2^G19V^* on Cln2 protein expression. Thus, the data suggest that ras2^G19V^ disrupts the mechanisms by which Cln3 induces transcription of late G1 phase cyclins, which is a critical step in the molecular mechanisms that initiate entry into the cell cycle.

### *ras2^G19V^* is unlikely to control G1 cyclin expression via a simple PKA-Whi3 signaling axis

Previous studies suggested that *ras2^G19V^*could influence G1 cyclin expression and cell cycle entry via a signaling axis in which Ras activates PKA to inhibit Whi3, which is thought to bind and inhibit expression of G1 cyclin mRNAs (Garí et al., 2001; Mizunuma et al., 2013; Cai and Futcher, 2013). We tested this model by comparing the effects of *ras2^G19V^*to the effects of *whi3Δ* or inhibition of PKA. Expression of *ras2^G19V^* and *whi3Δ* both caused an increase in Cln3 protein levels and a *lexA-ras2^G19V^ whi3*Δ double mutant did not show additive effects on Cln3 protein levels. These observations are consistent with a model in which *ras2^G19V^* drives an increase in Cln3 protein levels via inhibition of Whi3 but do not rule out alternative models. However, the data do not support the idea that *ras2^G19V^* influences Cln2 protein levels via a PKA-Whi3 signaling axis. For example, *ras2^G19V^* and inhibition of PKA caused similar effects on Cln2 protein levels, which would not be expected if *ras2^G19V^*drives a decrease in Cln2 protein levels via hyperactivation of PKA. Overall, the data are most consistent with a model in which hyperactive signaling from *ras2^G19V^*influences Cln2 protein levels via a PKA-independent pathway. Testing this model will require additional work to define the signaling steps by which *ras2^G19V^*blocks normal expression of Cln2.

### The effects of *ras2^G19V^* on late G1 cyclin expression are dependent upon Whi5

Previous studies suggested that Cln3 initiates transcription of late G1 phase cyclins at least partly via inhibition of Whi5 (de Bruin et al., 2004; Costanzo et al., 2004). Here, we found that expression of *ras2^G19V^* does not delay expression of Cln2 in *whi5Δ* cells, which suggests that *ras2^G19V^* may disrupt the mechanism by which Cln3 inactivates Whi5. Previous studies suggested that a Cln3/Cdk1 complex directly phosphorylates and inactivates Whi5; however, more recent work has definitively shown that Cln3 is not required for phosphorylation of Whi5 during cell cycle entry (Kõivomägi et al., 2021; Bhaduri et al., 2015). Thus, the mechanisms that inactivate Whi5 are poorly understood.

Whi5 appears to be functionally similar to Rb in mammalian cells, as both are repressors of late G1 phase cyclin transcription. However, Whi5 and Rb show no sequence homology and recent evidence suggests that they may be regulated via different mechanisms (Kõivomägi et al., 2021; Bhaduri et al., 2015). It has therefore remained unclear whether the mechanisms that control Whi5 and Rb are conserved. Further investigation of the mechanisms by which *ras2^G19V^* influences Whi5 activity could therefore lead to a better understanding of the relationship between signals that control Whi5 and Rb.

### Evidence for Cln2-dependent negative regulation of Cln3

Expression of Cln2 protein was reduced and delayed by *ras2^G19V^* and also by inhibition of PKA. In both contexts, the window of Cln3 protein expression was prolonged, which could be explained by a model in which Cln2 represses expression of Cln3. Consistent with this, we found that overexpression of Cln2 causes reduced expression of Cln3 protein, while loss of Cln1/2 appeared to increase and prolong expression of Cln3. Previous studies have shown that this kind of feedback regulation works at other times during the cell cycle in yeast (Amon et al., 1993). The discovery that Cln3 appears to be regulated by Cln2-dependent negative feedback suggests a new entry point for further exploration of the mechanisms that control cell cycle entry.

### Analysis of aberrant Ras signaling in yeast may provide new insight into cancer cell biology

Figure 10 shows a simplified model of the mechanisms that are thought to control cell cycle entry, in which the steps that appear to be influenced by *ras2^G19V^*are highlighted. Our discovery that *ras2^G19V^* influences key steps in cell cycle entry, potentially via PKA-independent mechanisms, provides new insight into how *ras2^G19V^* influences cell size and cell cycle progression. In mammalian cells, there is evidence that hyperactive Ras influences expression of cyclin D via mechanisms that are independent of the canonical MAP kinase signaling pathway that has been thought to mediate many functions of Ras (Pruitt et al., 2000; Pruitt and Der, 2001; Kerkhoff and Rapp, 1998; Coleman et al., 2004; Muise-Helmericks et al., 1998). Together, these observations indicate that much remains to be learned about how aberrant Ras signaling influences basic cell biology. Further investigation of the mechanisms by which hyperactive Ras influences cell size and the cell cycle in yeast could yield broadly relevant insights into basic cell biology as well as cancer cell biology.

**Figure 10.**
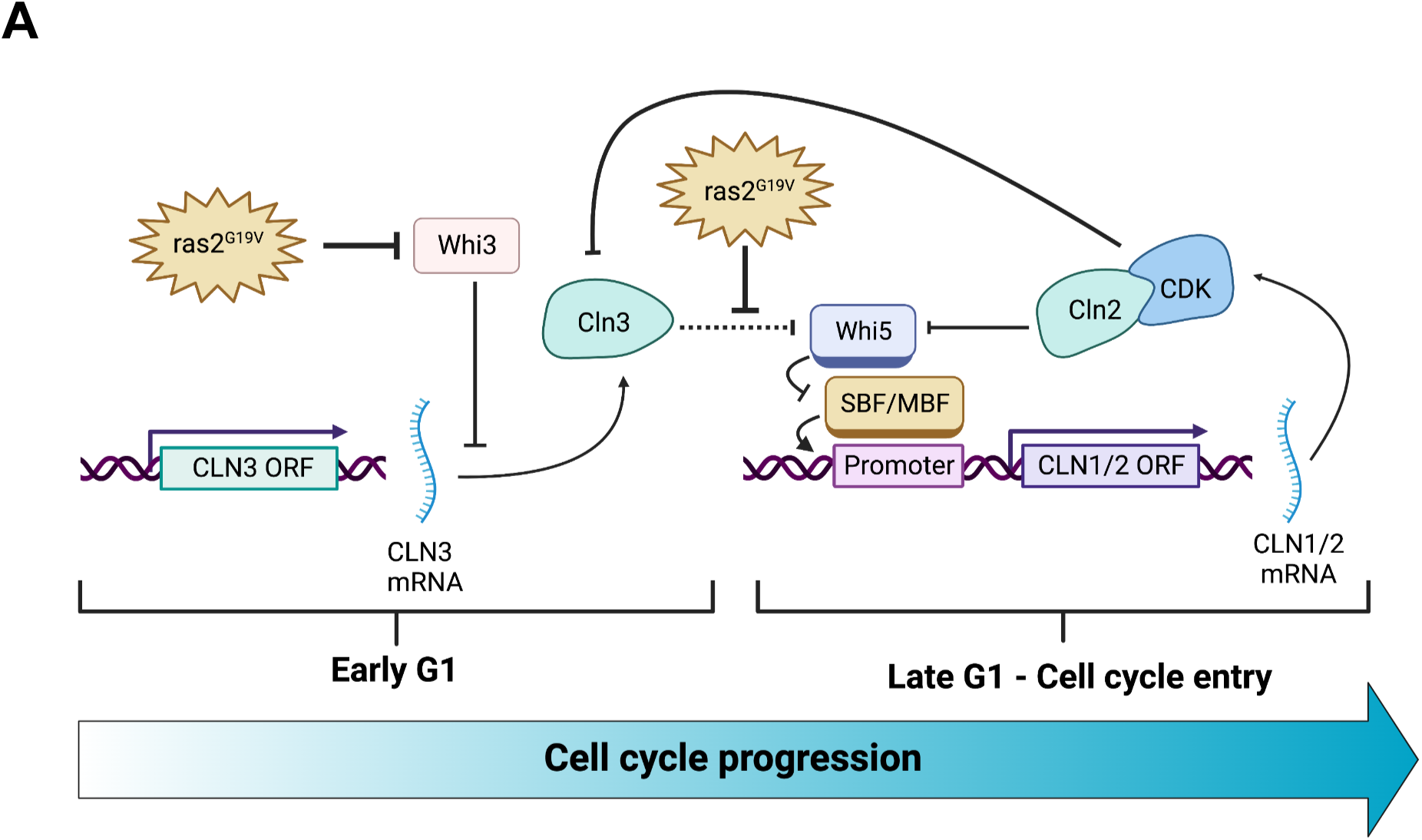
Proposed model. A simplified model of the steps that control cell cycle entry in which events that appear to be influenced by *ras2^G19V^* are highlighted.

## Materials and Methods

### Yeast strains and media

The genotypes of the strains used in this study are listed in Table 1. All strains are in the W303 background (*leu2-3,112 ura3-1 can1-100 ade2-1 his3-11,15 trp1-1 GAL+, ssd1-d2*). Genetic alterations, such as addition of epitope tags, promoter swaps, and gene deletions were carried out using homologous recombination at the endogenous locus (Longtine et al., 1998; Janke et al., 2004). Strains that express *RAS2* or *ras2^G19V^* from the estradiol-inducible lexA promoter at the *RAS2* locus were generated by homologous recombination as previously described (Ottoz et al., 2014). Briefly, *lexA* promoter elements were amplified and integrated upstream of wild type *RAS2* locus (oligos: *TAACCGTTTTCGAATTGAAAGGAGATATACAGAAAAAAAACGAGAGCTTGCC-TTGTCCCC* and *GTACTCTCTTATGTTCGACTTGTTCAAAGGCATAAGCTTGATATCGA-ATTCCTG*). The *ras2^G19V^* mutation was incorporated into the 3’ oligo (oligos: *TAACC-GTTTTCGAATTGAAAGGAGATATACAGAAAAAAAACGAGAGCTTGCCTTGTCCCC, ATTGGGTCAATTGTATGGTCAAAGCAGATTTACCAACACCAACACCACCAACGACGACTAGCTTGTACTCTCTTATGTTCGACTTGTTCAAAGGCATAAGCTTGATATCGAATTCCTG*). The *ras2^G19V^* mutant was verified by Sequencing.

**Table 1.** List of budding yeast strains used in this study.

| **Strain** | **Genotype** | **Source** | **Figures** |
| --- | --- | --- | --- |
| **DK4040** | *MATa bar1 ras2Δ::URA3* | This study | Figure 1 |
| **DK4042** | *MATa bar1 ras1Δ::KANMX* | This study | Figure 1 and Figure 1–figure supplement 1 |
| **DK4064** | *MATa bar1 ras2<sup>G19V</sup>* | This study | Figure 1 and Figure 1–figure supplement 1 |
| **DK4099** | *MATa bar1 ras1Δ::KANMX ras2<sup>G19V</sup>* | This study | Figure 1–figure supplement 1 |
| **DK4333** | *MATa bar1 ira2ΔirKANMX* | This study | Figure 1 |
| **DK4426** | *MATa bar1 LexA-ER-B112::HIS3* | This study | Figure 2, Figure 3, Figure 5 |
| **DK4450** | *MATa bar1 LexA-ER-B112::HIS3 HPHNT1::(2x)lexA-RAS2* | This study | Figure 2, Figure 3 |
| **DK4452** | *MATa bar1 LexA-ER-B112::HIS3 HPHNT1::(2x)lexA-ras2<sup>G19V</sup>* | This study | Figure 2, Figure 3, Figure 5 |
| **DK4495** | *MATa bar1 LexA-ER-B112::HIS3 CLN3-6xHA::NATNT2 CLN2-13xMYC::KANMX* | This study | Figure 4, Figure 4–figure supplement 1, Figure 7, Figure 7–figure supplement 1, and Figure 8 |
| **DK4496** | *MATa bar1 LexA-ER-B112::HIS3 HPHNT1::(2x)lexA-RAS2 CLN3-6xHA::NATNT2 CLN2-13xMYC::KANMX* | This study | Figure 4, Figure 4–figure supplement 1 |
| **DK4497** | *MATa bar1 LexA-ER-B112::HIS3 HPHNT1::(2x)lexA-ras2<sup>G19V</sup> CLN3-6xHA::NATNT2 CLN2-13xMYC::KANMX* | This study | Figure 4, Figure 4–figure supplement 1, Figure 7, Figure 7–figure supplement 1, and Figure 8 |
| **DK4518** | *MATa bar1 CLN3-6xHA::NATNT2 CLN2-13xMYC::KANMX whi3Δ::URA3* | This study | Figure 7, Figure 7–figure supplement 1 |
| **DK4519** | *MATa , bar1, LexA-ER-B112::HIS3, HPHNT1::(2x)lexA-ras2<sup>G19V</sup> CLN3-* | This study | Figure 7, Figure 7–figure supplement 1 |
|  | *6xHA::NATNT2, CLN2-13xMYC::KANMX whi3Δ::URA3* |  |  |
| **DK4580** | *MATa bar1 LexA-ER-B112::HIS3 HPHNT1::(2x) lexA-ras2<sup>G19V</sup> CLN2-9xMYC::NATNT2* | This study | Figure 5 |
| **DK4589** | *MATa bar1 CLN3-6xHA::HIS3 KANMX::Gal-CLN2* | This study | Figure 9 |
| **DK4597** | *MATa bar1 LexA-ER-B112::HIS3 HPHNT1::(2x) lexA-ras2<sup>G19V</sup> KANMX::MET25-CLN2-9xMYC::NATNT2* | This study | Figure 5 |
| **DK4629** | *MATa bar1 cln1ΔclTRP cln2Δ::LEU CLN3-6xHA::HIS3* | This study | Figure 9 |
| **DK4647** | *MATa CLN3-6xHA::NATNT2 CLN2-13xMyc::KANMX* | This study | Figure 6 |
| **DK4648** | *MATa CLN3-6xHA::NATNT2 CLN2-13xMyc::KANMX tpk1(M147G) tpk2 (M164G) tpk3 (M165G)* | This study | Figure 6 |
| **DK4731** | *MATa bar1 LexA-ER-B112::HIS3 HPHNT1::(2x) lexA-ras2<sup>G19V</sup> CLN3-6xHA::NATNT2 CLN2-13xMYC::KANMX whi5Δ::URA3* | This study | Figure 8 |

Cells were grown in YP medium (1% yeast extract, 2% peptone) that contained 40 mg/L adenine and a carbon source. Rich carbon medium (YPD) contained 2% dextrose, while poor carbon medium (YPG/E) contained 2% glycerol and 2% ethanol. In experiments using the ATP analog inhibitors 1-NM-PP1 or 3-MOB-PP1 no additional adenine was added to the media. All ATP analog inhibitors were solubilized in 100% DMSO. 3-MOB-PP1 was a gift from Kevin Shokat (UCSF). β-estradiol (#50-28-2 from Acros) was added to cultures from a 10 mM stock in 100% ethanol.

### Cell cycle time-courses and serial dilution assays

Cell cycle time courses were carried out as previously described (Harvey et al., 2011). Briefly, cells were grown to log phase at room temperature overnight in YPD or YPG/E medium to an optical density (OD^600^) of 0.5 - 0.7. Cultures were adjusted to the same optical density and were then arrested in G1 phase with alpha factor at room temperature for 3 hours. *bar1* strains were arrested with 0.5 µg/mL alpha factor, while *BAR1*+ strains were arrested with 15 µg/mL alpha factor. Cells were released from the arrest by washing 3 times with YPD or YPG/E. All time courses were carried out at 30°C unless otherwise noted, and alpha factor was added back at 40 minutes to prevent initiation of a second cell cycle. For experiments involving induced expression of *RAS2* or *ras2^G19V^*, cells were arrested for 3 hours and β-estradiol (200 nM) was added 1.5 hours before release. Cells were released from the arrest by washing 3 times with fresh YPD containing 200 nM β-estradiol.

Serial dilution assays were carried out by growing cells overnight in YPD to an OD^600^ of 0.4. A series of 10-fold dilutions were prepared, spotted on YPD plates, and grown for 2 days at 30°C.

### Northern blotting

Gel-purified PCR products were used to generate radio-labeled probes to detect *CLN2* and *ACT1* mRNAs by Northern blotting (oligonucleotides: TCAAGTTGGATGCAATTTGCAG, *TGAACCAATGATCAATGATTACGT*; *ACT1* oligonucleotides: TCATACCTTCTACAACGA-ATTGAGA and ACACTTCATGATGGAGTTGTAAGT. Probes were labeled using the Megaprime DNA labelling kits (GE Healthcare). Northern blotting was carried out as previously described (Cross and Tinkelenberg, 1991; Kellogg and Murray, 1995). *CLN3* blots were stripped and reprobed for *ACT1* to control for loading.

### ChIP and qPCR

ChIPs were performed as previously described (Voth et al., 2007). Yeast cells were collected at an OD of 0.6–0.8 and cross-linked in 1% formaldehyde for 20 min at room temperature. Cross-linking was quenched with 0.125 M glycine for 5 min, and cells were washed twice with 1X TBS (0.05 M Tris, 0.15 M NaCl, pH 7.6). Cell pellets were resuspended in lysis buffer (0.1% deoxycholic acid, 1 mM EDTA, 50 mM Hepes, pH 7.5, 140 mM NaCl, 1% Triton X-100 supplemented with protease inhibitors) and were lysed with 0.5-mm glass beads (BioSpec #11079105) in a cell disrupter (Mini-Beadbeater; BioSpec). After centrifugation, the pellet was washed with lysis buffer and sonicated to a shearing size of <500 nucleotides using a bath sonicator (Biorupter XL; Diagenode). The sonicated material was centrifuged, and the supernatant was collected for immunoprecipitation.

Chromatin immunoprecipitations were performed overnight at 4°C using 500–700 µg of chromatin and a mouse monoclonal anti-HA antibody (12CA5, Gift of David Toczyski, University of California, San Francisco) bound to Protein A Dynabeads (Thermo Fisher #10001D). The beads were washed twice with lysis buffer (50 mM HEPES-KOH, pH 7.5, 140mM NaCl, 1mM EDTA, 1% Triton X-100, 0.1% Sodium Deoxycholate), twice with high salt buffer (lysis buffer with 500 mM NaCl), twice with LiCl buffer (0.5% deoxycholic acid, 1 mM EDTA, 250 mM LiCl, 0.5% NP-40, 10 mM Tris-HCl, pH 8.0), and once with TE buffer (10 mM Tris-HCl, pH 8.0, 1 mM EDTA). Cross-links were reversed overnight in elution buffer (5 mM EDTA, 0.5% SDS, 0.3 mM NaCl, 10 mM Tris-HCl, pH 8.0) at 65°C. DNA was purified using the ChIP DNA Clean & Concentrator purification kit (Zymo #D5201). Quantitative PCR reactions were performed using a detection system (LightCycler480 II; Roche). A standard curve representing a range of concentrations of input samples was used for quantifying the amount of product for each sample with each primer set. All ChIP samples were normalized to corresponding input control samples, to a genomic reference region on chromosome I, and to a genetically identical untagged strain as a control. (ChIP primers for the *CLN2* promoter region: *CAATTCATGCGCGCTTTACC, TCTTCGCTAGGTATCCGCAT.* ChIP primers for the chromosome I control region: *GTTTATAGCGGGCATTATGCGTAGATCAG* and *GTTCCTCTAGAATTTTTCCACTCGCACA-TT*.)

### Western blotting and quantification

For western blotting, 1.6 ml samples were taken from cultures and pelleted in a microfuge at 13,200 rpm for 30 sec before aspirating the supernatant and adding 250 µL of glass beads and freezing on liquid nitrogen. Cells were lysed in 140 µL of 1X SDS-PAGE sample buffer (65 mM Tris-HCl, pH 6.8, 3% SDS, 10% glycerol, 100 mM βglycerophosphate, 50 mM NaF, 5% β-mercaptoethanol, 2 mM PMSF, and bromophenol blue) by bead beating in a Biospec Mini-Beadbeater-16 at 4°C for two minutes. The lysate was centrifuged for 15 seconds to bring the sample to the bottom of the tube and was then incubated in a 100°C water bath for 5 minutes followed by a centrifugation for five minutes at 13,200 rpm. Lysates were loaded onto 10% acrylamide SDS-PAGE gels that were run at a constant current setting of 20 mA per gel at 165 V. Gels were transferred to nitrocellulose membrane in a BioRad Trans-Blot Turbo Transfer system. Blots were probed overnight at 4°C in 4% milk in western wash buffer (1x PBS + 250 mM NaCl + 0.1% Tween-20) with mouse monoclonal anti-HA antibody (12CA5, Gift of David Toczyski, University of California, San Francisco), mouse monoclonal anti-Myc (2276S from Cell Signaling), polyclonal anti-Clb2 antibody, polyclonal anti-Nap1 antibody, polyclonal anti-Ypk1 antibody, or polyclonal rabbit anti-T662P antibody (Gift from Ted Powers, University of California, Davis). Western blots using anti-T662P antibody were first blocked using TBST (10 mM Tris-Cl, pH 7.5, 100 mM NaCl, and 0.1% Tween-20) + 4% milk, followed by one wash with TBST, then overnight incubation with anti-T662P antibody in TBST + 4% BSA. Western blots were incubated in secondary donkey anti-mouse (GE Healthcare NA934V) or donkey anti-rabbit (GE Healthcare NXA931 or Jackson Immunoresearch 711-035-152) antibody conjugated to HRP at room temperature for 60-90 min before imaging with Advansta ECL chemiluminescence reagents in a BioRad ChemiDoc imaging system. Western blots were quantified using BioRad Imagelab software v6.0.1. Relative signal was calculated by normalizing to a loading control and then setting all other samples to a reference of either the zero-minute time point for time-course experiments or to wild type for log-phase comparisons (see figure legends for details).

### Cell size analysis by Coulter Channelizer and bud emergence

Yeast cells were grown overnight at 22°C to mid-log phase (OD600 less than 0.7). Cells were fixed with 3.7% formaldehyde for 30 min and were then washed with PBS + 0.02% Tween-20 + 0.1% sodium azide before measuring cell size using a Z2 Coulter Channelyzer as previously described (Lucena et al. 2018) using Z2 AccuComp v3.01a software. For log phase cultures, each cell size plot is an average of three independent biological replicates in which each biological replicate is the average of two technical replicates. The percentage of budded cells was calculated by counting >200 cells at each time point using a Zeiss Axioskop 2 (Carl Zeiss).

### Experimental replicates and Statistical analysis

All experiments were repeated for a minimum of 3 biological replicates. Biological replicates are defined as experiments that are carried out on different days with different cultures. Figures present data from representative biological replicates and Coulter Counter data represent the average of biological replicates. For the statistical analyses, one-tailed unpaired t test was performed using Prism 9 (Grahpad). p values are described in each figure legend.

### Data availability

All strains and reagents are available upon request. The authors affirm that all data necessary for confirming the conclusions of the article are present within the article, figures, and tables.

---

**Figure 1–figure supplement 1.**
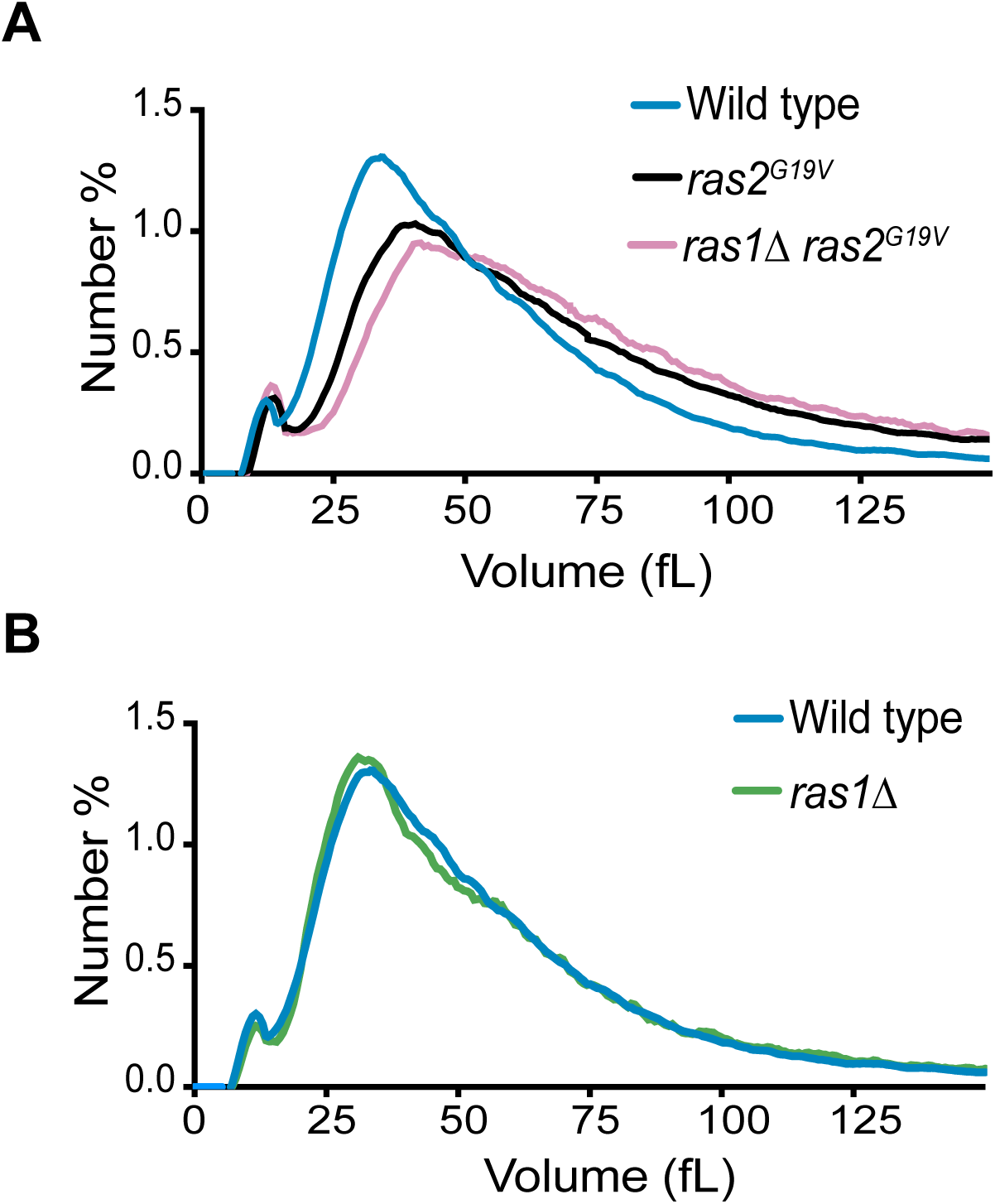
(A-B) Yeast cells were grown to log phase in YPD and cell size was measured using a Coulter counter. (A) Coulter counter size plots comparing Wild type, *ras2^G19V^,* and *ras1Δ ras2^G19V^* cells. (B) Coulter counter size plots comparing wild type, and *ras1Δ* cells.

**Figure 4–figure supplement 1.**
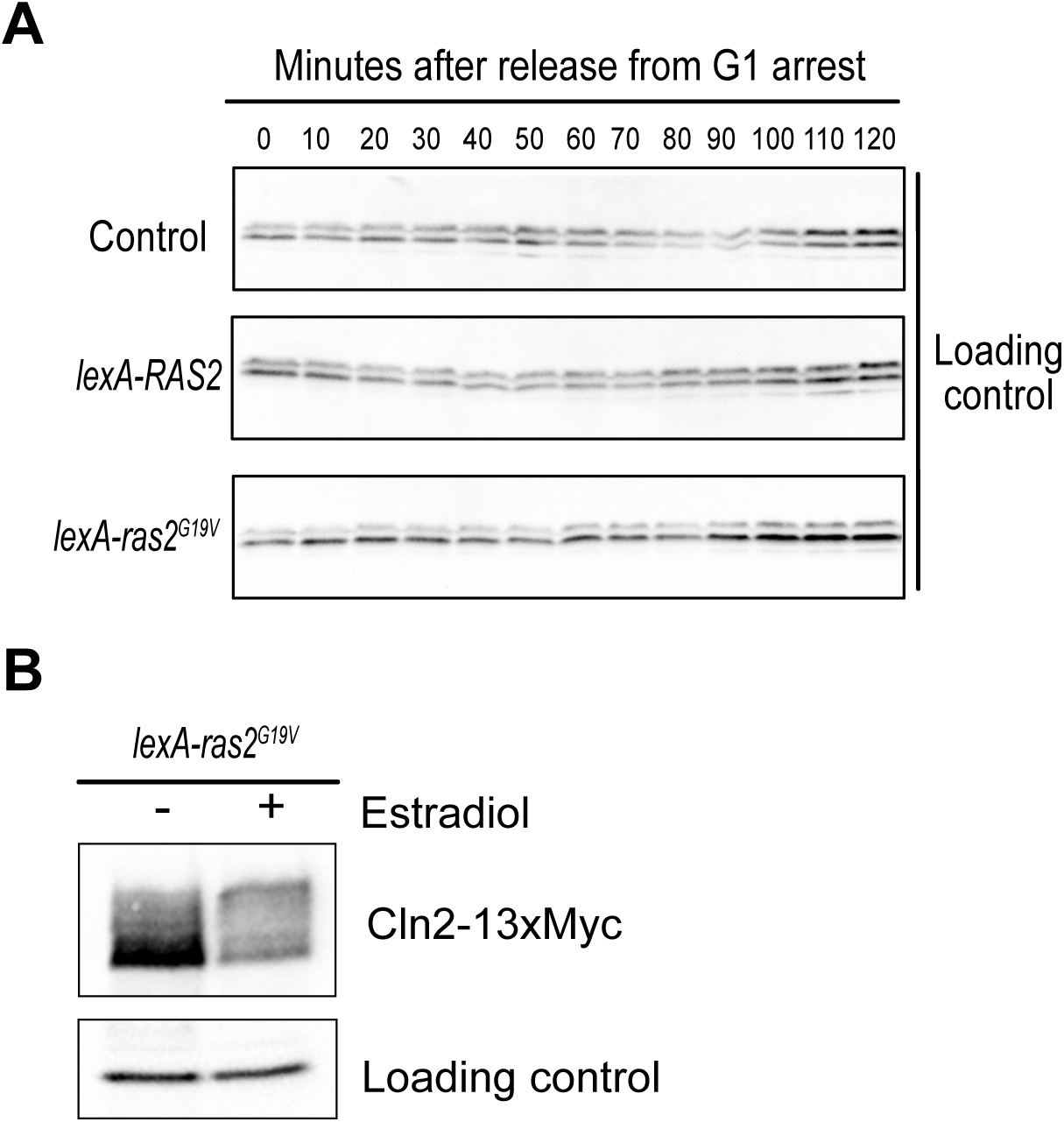
(A) Loading controls for the data in Figures 4 A,B. Levels of Nap1 protein were analyzed for a loading control. (B) Cells were grown to log-phase overnight in YPD and then treated with vehicle or 200 nM estradiol for 1.5 hours. The levels of Cln2-13xMyc protein were analyzed by western blot.

**Figure 7–figure supplement 1.**
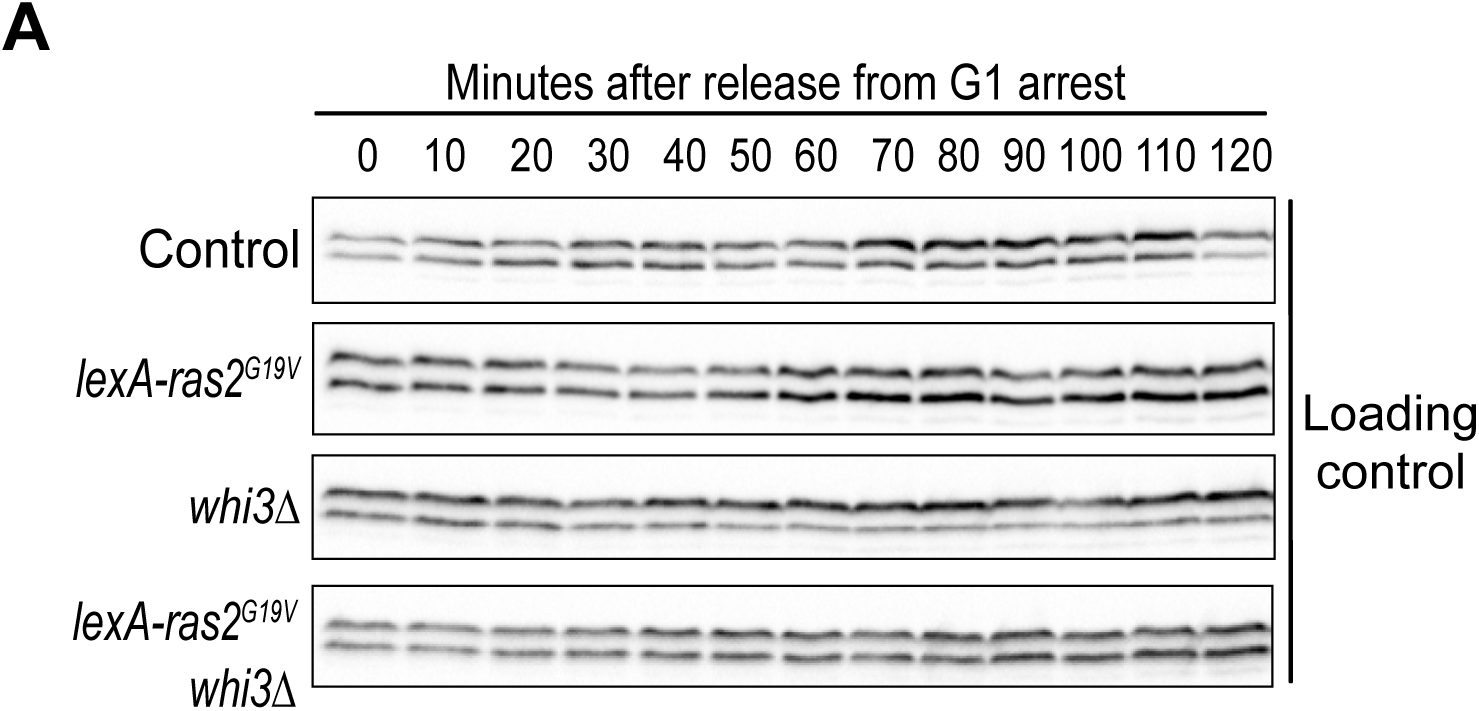
Loading controls for the data in Figures 7C,D. Levels of Nap1 protein were analyzed for a loading control.

## References

1. Amon, A., M. Tyers, B. Futcher, and K. Nasmyth. 1993. Mechanisms that help the yeast cell cycle clock tick: G2 cyclins transcriptionally activate G2 cyclins and repress G1 cyclins. Cell. 74:993–1007. doi:10.1016/0092-8674(93)90722-3.

2. Asa, S.L. 2019. The Current Histologic Classification of Thyroid Cancer. Endocrinology and Metabolism Clinics of North America. 48:1–22. doi:10.1016/j.ecl.2018.10.001.

3. Asadullah, S. Kumar, N. Saxena, M. Sarkar, A. Barai, and S. Sen. 2021. Combined heterogeneity in cell size and deformability promotes cancer invasiveness. Journal of Cell Science. 134:jcs250225. doi:10.1242/jcs.250225.

4. Baroni, M.D., E. Martegani, P. Monti, and L. Alberghina. 1989. Cell Size Modulation by CDC25 and RAS2 Genes in Saccharomyces cerevisiae. MOL. CELL. BIOL. 9. doi:10.1128/mcb.9.6.2715-2723.1989.

5. Baroni, M.D., P. Monti, and L. Alberghina. 1994. Repression of growth-regulated Gl cyclin expression by cyclic AMP in budding yeast. Nature. 371:339–342. doi:10.1038/371339a0.

6. Bhaduri, S., E. Valk, M.J. Winters, B. Gruessner, M. Loog, and P.M. Pryciak. 2015. A Docking Interface in the Cyclin Cln2 Promotes Multi-site Phosphorylation of Substrates and Timely Cell-Cycle Entry. Current Biology. 25:316–325. doi:10.1016/j.cub.2014.11.069.

7. Brimo, F., R. Montironi, L. Egevad, A. Erbersdobler, D.W. Lin, J.B. Nelson, M.A. Rubin, T. van der Kwast, M. Amin, and J.I. Epstein. 2013. Contemporary Grading for Prostate Cancer: Implications for Patient Care. European Urology. 63:892–901. doi:10.1016/j.eururo.2012.10.015.

8. Broek, D., T. Toda, T. Michaeli, L. Levin, M. Zoller, S. Powers, C. Birchmeier, and M. Wigler. 1987. The S. Cerevisiae Cdc25 Gene Product Regulates the Ras/Adenylate Cyclase Pathway. Cell Press. 48:11. doi:10.1016/0092-8674(87)90076-6.

9. de Bruin, R.A.M., W.H. McDonald, T.I. Kalashnikova, J. Yates, and C. Wittenberg. 2004. Cln3 Activates G1-Specific Transcription via Phosphorylation of the SBF Bound Repressor Whi5. Cell. 117:887–898. doi:10.1016/j.cell.2004.05.025.

10. Cai, Y., and B. Futcher. 2013. Effects of the Yeast RNA-Binding Protein Whi3 on the Half-Life and Abundance of CLN3 mRNA and Other Targets. PLoS ONE. 8:e84630. doi:10.1371/journal.pone.0084630.

11. Cazzanelli, G., F. Pereira, S. Alves, R. Francisco, L. Azevedo, P. Dias Carvalho, A. Almeida, M. Côrte-Real, M. Oliveira, C. Lucas, M. Sousa, and A. Preto. 2018. The Yeast Saccharomyces cerevisiae as a Model for Understanding RAS Proteins and their Role in Human Tumorigenesis. Cells. 7:14. doi:10.3390/cells7020014.

12. Coleman, M.L., C.J. Marshall, and M.F. Olson. 2004. RAS and RHO GTPases in G1-phase cell-cycle regulation. Nat Rev Mol Cell Biol. 5:355–366. doi:10.1038/nrm1365.

13. Costanzo, M., J.L. Nishikawa, X. Tang, J.S. Millman, O. Schub, K. Breitkreuz, D. Dewar, I. Rupes, B. Andrews, and M. Tyers. 2004. CDK Activity Antagonizes Whi5, an Inhibitor of G1/S Transcription in Yeast. Cell. 117:899–913. doi:10.1016/j.cell.2004.05.024.

14. Cross, F.R. 1988. DAFJ, a Mutant Gene Affecting Size Control, Pheromone Arrest, and Cell Cycle Kinetics of Saccharomyces cerevisiae. MOL. CELL. BIOL. 8:10. doi:10.1128/mcb.8.11.4675-4684.1988.

15. Dirick, L., T. Böhm, and K. Nasmyth. 1995. Roles and regulation of Cln-Cdc28 kinases at the start of the cell cycle of Saccharomyces cerevisiae. The EMBO Journal. 14:4803–4813. doi:10.1002/j.1460-2075.1995.tb00162.x.

16. Garí, E., T. Volpe, H. Wang, C. Gallego, B. Futcher, and M. Aldea. 2001. Whi3 binds the mRNA of the G _1_ cyclin *CLN3* to modulate cell fate in budding yeast. Genes Dev. 15:2803– 2808. doi:10.1101/gad.203501.

17. Gothwal, M., A. Nalwa, P. Singh, G. Yadav, M. Bhati, and N. Samriya. 2021. Role of Cervical Cancer Biomarkers p16 and Ki67 in Abnormal Cervical Cytological Smear. J Obstet Gynecol India. 71:72–77. doi:10.1007/s13224-020-01380-y.

18. Guerra, P., L.-A.P.E. Vuillemenot, Y.B. van Oppen, M. Been, and A. Milias-Argeitis. 2022. TORC1 and PKA activity towards ribosome biogenesis oscillates in synchrony with the budding yeast cell cycle. Journal of Cell Science. 135:jcs260378. doi:10.1242/jcs.260378.

19. Hall, D.D. 1998. Regulation of the Cln3-Cdc28 kinase by cAMP in Saccharomyces cerevisiae. The EMBO Journal. 17:4370–4378. doi:10.1093/emboj/17.15.4370.

20. Harvey, S.L., G. Enciso, N. Dephoure, S.P. Gygi, J. Gunawardena, and D.R. Kellogg. 2011. A phosphatase threshold sets the level of Cdk1 activity in early mitosis in budding yeast. MBoC. 22:3595–3608. doi:10.1091/mbc.e11-04-0340.

21. Hobbs, G.A., C.J. Der, and K.L. Rossman. 2016. RAS isoforms and mutations in cancer at a glance. Journal of Cell Science. jcs.182873. doi:10.1242/jcs.182873.

22. Hoda, R.S., R. Lu, R.N. Arpin, M.W. Rosenbaum, and M.B. Pitman. 2018. Risk of malignancy in pancreatic cysts with cytology of high-grade epithelial atypia. Cancer Cytopathology. 126:773–781. doi:10.1002/cncy.22035.

23. Hubler, L., J. Bradshaw-Rouse, and W. Heideman. 1993. Connections between the Ras-cyclic AMP pathway and G1 cyclin expression in the budding yeast Saccharomyces cerevisiae. Mol. Cell. Biol. 13:6274–6282. doi:10.1128/MCB.13.10.6274.

24. Jorgensen, P., J.L. Nishikawa, B.-J. Breitkreutz, and M. Tyers. 2002. Systematic Identification of Pathways That Couple Cell Growth and Division in Yeast. Science. 297:395–400. doi:10.1126/science.1070850.

25. Jorgensen, P., and M. Tyers. 2004. How Cells Coordinate Growth and Division. Current Biology. 14:R1014–R1027. doi:10.1016/j.cub.2004.11.027.

26. Kellogg, D.R., and P.A. Levin. 2022. Nutrient availability as an arbiter of cell size. Trends in Cell Biology. S0962892422001477. doi:10.1016/j.tcb.2022.06.008.

27. Kerkhoff, E., and U.R. Rapp. 1998. Cell cycle targets of Ras/Raf signalling. Oncogene. 17:1457–1462. doi:10.1038/sj.onc.1202185.

28. Kõivomägi, M., M.P. Swaffer, J.J. Turner, G. Marinov, and J.M. Skotheim. 2021. G _1_ cyclin–Cdk promotes cell cycle entry through localized phosphorylation of RNA polymerase II. Science. 374:347–351. doi:10.1126/science.aba5186.

29. Landry, B.D., J.P. Doyle, D.P. Toczyski, and J.A. Benanti. 2012. F-Box Protein Specificity for G1 Cyclins Is Dictated by Subcellular Localization. PLoS Genet. 8:e1002851. doi:10.1371/journal.pgen.1002851.

30. Liu, S., C. Tan, M. Tyers, A. Zetterberg, and R. Kafri. 2022. What programs the size of animal cells? Front. Cell Dev. Biol. 10:949382. doi:10.3389/fcell.2022.949382.

31. Lucena, R., M. Alcaide-Gavilán, K. Schubert, M. He, M.G. Domnauer, C. Marquer, C. Klose, M.A. Surma, and D.R. Kellogg. 2018. Cell Size and Growth Rate Are Modulated by TORC2-Dependent Signals. Current Biology. 28:196–210.e4. doi:10.1016/j.cub.2017.11.069.

32. Matsumoto, K., I. Uno, and T. Ishikawa. 1983. Control of cell division in Saccharomyces cerevisiae mutants defective in adenylate cyclase and cAMP-dependent protein kinase. Experimental Cell Research. 146:151–161. doi:10.1016/0014-4827(83)90333-6.

33. Mizunuma, M., R. Tsubakiyama, T. Ogawa, A. Shitamukai, Y. Kobayashi, T. Inai, K. Kume, and D. Hirata. 2013. Ras/cAMP-dependent Protein Kinase (PKA) Regulates Multiple Aspects of Cellular Events by Phosphorylating the Whi3 Cell Cycle Regulator in Budding Yeast. Journal of Biological Chemistry. 288:10558–10566. doi:10.1074/jbc.M112.402214.

34. Muise-Helmericks, R.C., H.L. Grimes, A. Bellacosa, S.E. Malstrom, P.N. Tsichlis, and N. Rosen. 1998. Cyclin D Expression Is Controlled Post-transcriptionally via a Phosphatidylinositol 3-Kinase/Akt-dependent Pathway. Journal of Biological Chemistry. 273:29864–29872. doi:10.1074/jbc.273.45.29864.

35. Nash, R., G. Tokiwa, S. Anand, K. Erickson, and A.B. Futcher. 1988. The WHI1+ gene of Saccharomyces cerevisiae tethers cell division to cell size and is a cyclin homolog. The EMBO Journal. 7:4335–4346. doi:10.1002/j.1460-2075.1988.tb03332.x.

36. Nash, R.S., T. Volpe, and B. Futcher. 2001. Isolation and characterization of WHI3, a size-control gene of Saccharomyces cerevisiae. Genetics. 157:1469–1480. doi:10.1093/genetics/157.4.1469.

37. Nasmyth, K., and L. Dirick. 1991. The Role of SW4 and SW16 in the Activity of Gl Cyclins in Yeast. Cell. 66:995–1013. doi:10.1016/0092-8674(91)90444-4.

38. Ottoz, D.S.M., F. Rudolf, and J. Stelling. 2014. Inducible, tightly regulated and growth condition-independent transcription factor in Saccharomyces cerevisiae. Nucleic Acids Research. 42:e130–e130. doi:10.1093/nar/gku616.

39. Pruitt, K., and C.J. Der. 2001. Ras and Rho regulation of the cell cycle and oncogenesis. Cancer Letters. 171:1–10. doi:10.1016/S0304-3835(01)00528-6.

40. Pruitt, K., R.G. Pestell, and C.J. Der. 2000. Ras Inactivation of the Retinoblastoma Pathway by Distinct Mechanisms in NIH 3T3 Fibroblast and RIE-1 Epithelial Cells. Journal of Biological Chemistry. 275:40916–40924. doi:10.1074/jbc.M006682200.

41. Robinson, L.C., J.B. Gibbs, M.S. Marshall, I.S. Sigal, and K. Tatchell. 1987. CDC25: A Component of the RAS-Adenylate Cyclase Pathway in Saccharomyces cvisa e. Science. 235:4. doi:10.1126/science.3547648.

42. Sandlin, C.W., S. Gu, J. Xu, C. Deshpande, M.D. Feldman, and M.C. Good. 2022. Epithelial cell size dysregulation in human lung adenocarcinoma. PLoS ONE. 17:e0274091. doi:10.1371/journal.pone.0274091.

43. Sommer, R.A., J.T. DeWitt, R. Tan, and D.R. Kellogg. 2021. Growth-dependent signals drive an increase in early G1 cyclin concentration to link cell cycle entry with cell growth. eLife. 10:e64364. doi:10.7554/eLife.64364.

44. Stuart, D., and C. Wittenberg. 1995. CLN3, not positive feedback, determ the timing of CLN2 transcription in cycling cells. Genes and Development. doi:10.1101/gad.9.22.2780.

45. Toda, T., D. Broek, T. Kataoka, I. Uno, T. Ishikawa, S. Powers, S. Cameron, J. Broach, K. Matsumoto, and M. Wigler. 1985. In yeast, RAS proteins are controlling elements of adenylate cyclase. Cell. 40:27–36. doi:10.1016/0092-8674(85)90305-8.

46. Toda, T., P. Sass, M. Zoller, J.D. Scott, B. McMULLEN, M. Hurwitz, E.G. Krebs, M. Wigler, and B. McMullen. 1987. Cloning and Characterization of BCY1, a Locus Encoding a Regulatory Subunit of the Cyclic AMP-Dependent Protein Kinase in Saccharomyces cerevisiae. MOL. CELL. BIOL. 7:7. doi:10.1128/mcb.7.4.1371-1377.1987.

47. Tokiwa, G., T. Volpe, M. Tyers, and B. Futcher. 1994. Inhibition of G1 cyclin activity by the Ras/cAMP pathway in yeast. Nature. 371:4. doi:10.1038/371342a0.

48. Turner, J.J., J.C. Ewald, and J.M. Skotheim. 2012. Cell size control in yeast. Curr Biol. 22:R350–359. doi:10.1016/j.cub.2012.02.041.

49. Tyers, M., G. Tokiwa’, and B. Futcher. 1993. Comparison of the Saccharomyces cerevisiae G1 cyclins: Cln3 may be an upstream activator of Cln1, Cln2 and other cyclins. The EMBO Journal. 12:14. doi:10.1002/j.1460-2075.1993.tb05845.x.

50. Wang, H., L.B. Carey, Y. Cai, H. Wijnen, and B. Futcher. 2009. Recruitment of Cln3 Cyclin to Promoters Controls Cell Cycle Entry via Histone Deacetylase and Other Targets. PLoS Biol. 7:e1000189. doi:10.1371/journal.pbio.1000189.

51. Wang, H., E. Garí, E. Vergés, C. Gallego, and M. Aldea. 2004. Recruitment of Cdc28 by Whi3 restricts nuclear accumulation of the G1 cyclin–Cdk complex to late G1. EMBO J. 23:180–190. doi:10.1038/sj.emboj.7600022.

52. Weiss, R.A. 2020. A perspective on the early days of RAS research. Cancer Metastasis Rev. 39:1023–1028. doi:10.1007/s10555-020-09919-1.

53. Zaman, S., S.I. Lippman, L. Schneper, N. Slonim, and J.R. Broach. 2009. Glucose regulates transcription in yeast through a network of signaling pathways. Mol Syst Biol. 5:245. doi:10.1038/msb.2009.2.

54. Zapata, J., N. Dephoure, T. MacDonough, Y. Yu, E.J. Parnell, M. Mooring, S.P. Gygi, D.J. Stillman, and D.R. Kellogg. 2014. PP2ARts1 is a master regulator of pathways that control cell size. Journal of Cell Biology. 204:359–376. doi:10.1083/jcb.201309119.

